# CCR7+ CD4 T Cell Immunosurveillance Disrupted in Chronic SIV-Induced Neuroinflammation in Rhesus Brain

**DOI:** 10.1101/2023.08.28.555037

**Authors:** Sonny R. Elizaldi, Chase E Hawes, Anil Verma, Ashok R. Dinasarapu, Yashavanth Shaan Lakshmanappa, Brent T Schlegel, Dhivyaa Rajasundaram, Jie Li, Blythe P Durbin-Johnson, Zhong-Min Ma, Danielle Beckman, Sean Ott, Jeffrey Lifson, John H. Morrison, Smita S. Iyer

## Abstract

CD4 T cells survey and maintain immune homeostasis in the brain, yet their differentiation states and functional capabilities remain unclear. Our approach, combining single-cell transcriptomic analysis, ATAC-seq, spatial transcriptomics, and flow cytometry, revealed a distinct subset of CCR7+ CD4 T cells resembling lymph node central memory (T_CM_) cells. We observed chromatin accessibility at the CCR7, CD28, and BCL-6 loci, defining molecular features of T_CM_. Brain CCR7+ CD4 T cells exhibited recall proliferation and interleukin-2 production ex vivo, showcasing their functional competence. We identified the skull bone marrow as a local niche for these cells alongside other CNS border tissues. Sequestering T_CM_ cells in lymph nodes using FTY720 led to reduced CCR7+ CD4 T cell frequencies in the cerebrospinal fluid, accompanied by increased monocyte levels and soluble markers indicating immune activation. In macaques chronically infected with SIVCL57 and experiencing viral rebound due to cessation of antiretroviral therapy, a decrease in brain CCR7+ CD4 T cells was observed, along with increased microglial activation and initiation of neurodegenerative pathways. Our findings highlight a role for CCR7+ CD4 T cells in CNS immune surveillance and their decline during chronic SIV-induced neuroinflammation highlights their responsiveness to neuroinflammatory processes.

**GRAPHICAL ABSTRACT:** 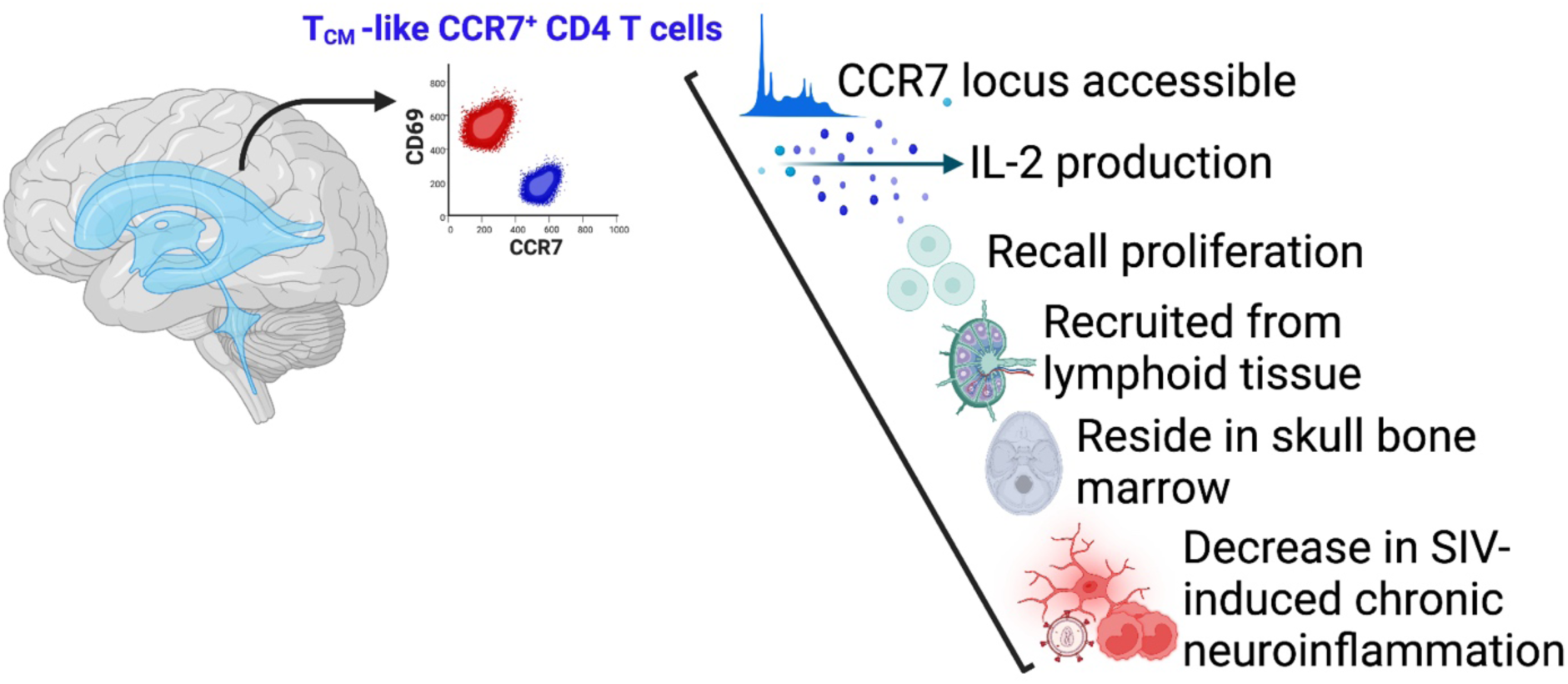

**In Brief:** Utilizing single-cell and spatial transcriptomics on adult rhesus brain, we uncover a unique CCR7+ CD4 T cell subset resembling central memory T cells (T_CM_) within brain and border tissues, including skull bone marrow. Our findings show decreased frequencies of this subset during SIV- induced chronic neuroinflammation, emphasizing responsiveness of CCR7+ CD4 T cells to CNS disruptions.

**Highlights:**

1. CCR7+ CD4 T cells survey border and parenchymal CNS compartments during homeostasis; reduced presence of CCR7+ CD4 T cells in cerebrospinal fluid leads to immune activation, implying a role in neuroimmune homeostasis.
2. CNS CCR7+ CD4 T cells exhibit phenotypic and functional features of central memory T cells (T_CM_) including production of interleukin 2 and the capacity for rapid recall proliferation. Furthermore, CCR7+ CD4 T cells reside in the skull bone marrow.
3. CCR7+ CD4 T cells are markedly decreased within the brain parenchyma during chronic viral neuroinflammation.

## INTRODUCTION

T lymphocytes play a critical role in immune protection against invading pathogens by surveying tissues. Upon infection, naive T cells encounter antigen in secondary lymphoid tissues undergo activation and proliferate. After pathogen clearance, a fraction of these activated T cells differentiate into diverse memory T cell subsets. Central memory T (T_CM_) cells ensure lasting protection by circulating through blood to secondary lymphoid organs. T_CM_ display high proliferative potential and contribute to effector and memory T cell (T_EM_) generation upon secondary antigen exposure (1, 2). Effector memory T cells (T_EM_) move between blood, spleen, and non-lymphoid tissues upon re-infection (3, 4). Tissue resident memory T (T_RM_) cells are stationed at non-lymphoid infection sites and, in the case of CD8 T cells, offer near sterilizing immunity (5). These distinct T memory subsets not only provide broad immune defense during infection and injury but also intricately modulate organ function during homeostasis.

The central nervous system (CNS) is recognized as an immunologically specialized site, where immune responses occur, although most lymphocytes are sequestered behind the CNS barriers during homeostasis (6, 7). Lymphocyte surveillance of the CNS primarily takes place at two barriers: the blood-cerebrospinal fluid barrier (BCSFB) and the blood-brain barrier (BBB). The CSF, generated at the choroid plexus, is among the main sites for T lymphocytes in the CNS during homeostasis and serves as a functional equivalent of lymph (6). The CSF is predominantly composed of T cells, which comprise 80% of cells in the CSF(8–11). CD4 T cells in the CSF express cell-surface markers consistent with T_CM_, such as CCR7, CD27, and CD45RO (8, 12). Beyond the CSF, recently, T_RM_ expressing phenotypic markers (CD69, CD103) have been identified in the human brain (13). Despite T cells occupying CNS niches during homeostasis, our understanding of specific T cell subsets establishing residency versus those surveilling the CNS and its border tissues remains incomplete. Bridging this gap is important to understand underpinnings of immune dysregulation during neuroinflammation.

We utilized the non-human primate model to define memory CD4 T cell subsets within the CNS. The rhesus macaque stands out as an exceptional model for investigating CNS T lymphocytes due to its resemblance to the human system, encompassing factors like genetic variability, neocortical structures, substantial dural and intricate leptomeningeal layers, and an array of memory T cell differentiation states. Utilizing techniques such as brain perfusion, which allows for the precise delineation of local brain immune populations while minimizing contamination from vasculature, studies in the macaque model offer a valuable platform for comprehensive immune analyses of distinct CNS compartments. Moreover, studying CNS memory T cells in macaques and humans presents distinct advantages over mouse models, as both have encountered pathogen-rich environments, fostering a diverse spectrum of memory T cell clones in lymphoid, peripheral, and CNS tissues.

Diverging from the conventional distribution patterns of memory T cells in non-lymphoid tissues, our study uncovers distinctive attributes of CD4 T cells within the brain parenchyma of rhesus macaques. These cells display elevated CD28 expression with clear division in CD69 and CCR7 expression, denoting T_RM_ and T_CM_ profiles, respectively. Notably, parenchymal CCR7+ CD4 T cells exhibit traits consistent with T_CM_, including interleukin 2 production, recall proliferation, and CD127 expression. Our analysis of single-cell transcriptomics identified T_CM_ cell clusters in the brain, and ATAC-seq data demonstrated chromatin accessibility of the pivotal lymph node homing gene, CCR7. Moreover, by sequestering T_CM_ cells in lymph nodes, we observed a decrease in CCR7+ CD4 T cell frequencies in the CSF, coupled with heightened monocyte levels and soluble markers indicating immune activation. Thus, the CNS harbors CD4 T cells primed for either immune surveillance or residency, based on CCR7 and CD69 expression. Enhancing our understanding of these distinct CNS CD4 T cell subsets during both homeostasis and infection will enable deep insights into their functions ultimately paving the way for innovative therapeutic strategies to combat neuroinflammatory disorders.

## RESULTS

### Single-cell transcriptomic analysis of CD45+ leukocytes identifies core T cell gene signatures in rhesus brain

In a previous study, we identified T cell-associated transcripts within synapse-dense brain regions using bulk RNA sequencing (10). However, due to scarcity of T cells within the brain and the abundance of neuronal and glial cell-associated transcripts, our ability to deeply characterize T cell heterogeneity and specific gene programs was limited. To bridge this gap, we employed single-cell transcriptomic analysis on cryopreserved CD45+ cells from rhesus brain tissue to elucidate transcriptional networks underlying memory T cell states in the non-inflamed brain parenchyma. T cells were easily distinguishable using flow cytometry, constituting an average of 20% of CD45+ cells with a CD4:CD8 ratio of 0.2:1 (**Figure S1**). Single-cell RNA sequencing (sc-RNA-seq) was performed on viably frozen CD45+ cells isolated from healthy rhesus macaque brains, and viable CD45+ splenocytes served as a lymphoid reference for T cell subset identification (**Figure 1A**). CD45+ cells were positively selected and enriched by flow sorting for purity and viability before sequencing (**Figures 1B-C**). A median of 4,952 CD45+ cells from the brain and 3,151 CD45+ cells from the spleen were sequenced per tissue sample, resulting in 19,000 single-cell transcriptomes that passed quality control (**Figures S2A-E**). Marker gene analysis validated our approach, demonstrating high expression of CD45 (PTPRC) in cells from both compartments (**Figure S2F**).

**Figure 1.**
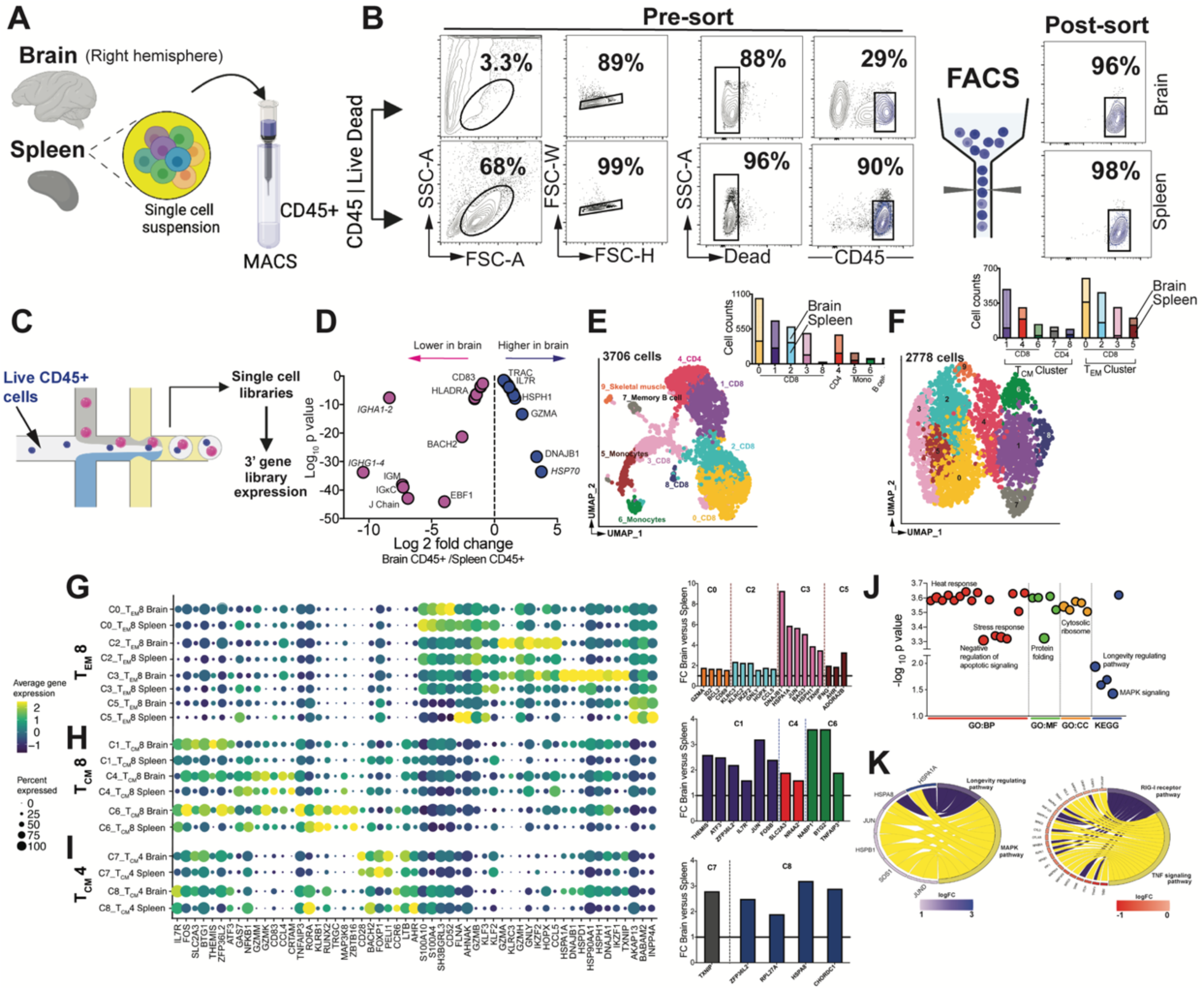
Single-cell transcriptomic analysis of CD45+ leukocytes identifies core T cell gene signatures in rhesus brain. (A-C) Schematic of single cell profiling on CD45+ cells in brain and spleen. **(D)** Differences in B and T cell gene abundance in brain versus spleen. **(E)** UMAP of scRNA-seq transcriptional profiles from brain and spleen shows 9 clusters. Cell clusters are color-coded based on cell identity assigned using Single R. Inset shows cell proportions in each cluster split by tissue type (bottom, spleen; top, brain) **(F)** UMAP shows 10 sub-clusters from T cell clusters in E. **(G-H)** Select marker genes of cell clusters. Dot size represents proportion of cells expressing gene and color designates expression level with bar graph representing genes significantly higher in brain relative to spleen for indicated clusters. **(I)** Chord plots show pathways and corresponding genes enriched versus underrepresented in T_CM_4 cell clusters.

Comparing transcriptomes between tissues, we observed distinct enrichment patterns. The spleen exhibited higher expression of B cell molecular signatures compared to the brain (**Figure 1D**). Notable enrichments included immunoglobulin-related genes (ENSMMUG00000015202 [human orthologs IGHG1-4], ENSMMUG00000002764 [human orthologs IGHA1 and IGHA2], IGHM, IGKC), genes linked to B cell function regulation (EBF1, BACH2, RelB), antigen presentation (CD74, HLA-DMB, HLA-DRA), and B cell markers (ALCAM, CD83, TRAF3)(14) (15) (16).This suggests a relatively lower B cell abundance in brain compared to spleen.

Conversely, CD45+ cells from the brain exhibited enriched T cell gene signatures. These included higher expression of transcripts associated with the T cell receptor α constant gene (TRAC), genes involved in TCR signaling (TAOK3, Sos1) (17) (18), genes regulating T cell metabolism (ERN1 and TXNIP(19), and genes linked to effector and central memory T cell programs (HSP70, DNAJB1, HSPH1, GZMA, NKG7, ID2, HELIOS (encoded by IKZF2), homeobox only protein (HOPX), and IL-7R (20) (21) (22). Additionally, the expression of the trafficking factor ITGβ2 suggested LFA-1 mediated T cell migration across the blood-brain barrier. Verification of B cell predominance in the spleen and T cell abundance in the brain was confirmed through established marker genes and cell type annotation of clusters (**Figures S2G-H**), supporting the presence of T cells within the rhesus macaque brain at homeostasis.

### Non-inflamed brain harbors both effector memory (T_EM_) resident memory (T_RM_) CD8 T cells

We next pursued high-resolution unsupervised clustering across tissue compartments. Through automated label transfer using blueprint_encode (23, 24), ten cell clusters were initially identified, and manual inspection along with marker gene analysis (25) confirmed four clusters as T cells (**Figures 1E, S3A**). To further dissect T cell subsets, the T cell clusters were isolated and independently reclustered, revealing 3 distinct T cell subtypes shared across the brain and spleen: Terminal effector memory CD8 T cells (T_EM_ 8 cluster; 0, 2, 3, 5), Central memory CD8 T cells (T_CM_ 8 cluster 1,4,6) and Central memory CD4 cells (T_CM_ 4 cluster 7 and 8) (**Figures 1F,S3B**).

Analysis of transcripts within the T_EM_8 cell clusters (C0, C2, C3, C5) unveiled distinct gene expression programs (**Figures 1G, S3C-D**) indicating functional diversity among brain CD8 T_EM_. T_EM_ 8 C0 exhibited elevated levels of transcripts consistent with *effector memory* cells, including Ca2+ binding proteins encoded by S100 family genes (S10010, S100A4) (26), SH3BGRL3 expressed by T_h_1 cells (27), regulatory receptor CD52 (28), molecules driving T cell activation (FLNA and the scaffold protein AHNAK (26, 29), cytolytic molecule GZMB, and transcription factors KLF2 and KLF3 (30, 31). Cells in T_EM_ 8 C2 were enriched for core *cytotoxic molecules* GZMA, GZMK, GZMB, GNLY, KLRC3, HELIOS, and HOPX expressed by human KIR+ CD8+ T cells (32) (33), and the β chemokine, CCL5. In contrast, cells in T_EM_ 8 C3 were enriched for DNAJ/Heatshock proteins regulating memory *T cell quiescence* (34, 35), Ikaros (encoded by IKZF1) and TXNIP, which suppress proliferation and inflammatory cytokines in T cells (36) (37). T_EM_ 8 C5 was enriched for genes regulating cell cycle progression and survival (AKAP13, BABAM2, INPP4A). Based on the higher relative expression of effector genes (GZMA, KLRC2-3, CCL5, IFNγ), residency and longevity genes (ID2, AHR, IKZF2, HOPX CD69, BCl2) in the brain compared to the spleen (**Bar graph in 1G**), we inferred that brain CD8 T_EM_ cell clusters encompass resident and effector memory subsets, consistent with the identification of CD8 T_EM_ and T_RM_ in the mouse and human brain (13, 38).

### Single-cell transcriptomic analysis reveals novel central memory CD4 and CD8 (T_CM_) subsets in brain

After investigating CD8 T_EM_ clusters, we shifted our focus to the remaining CD8 T cell clusters (C1, C4, C6) annotated as central memory (T_CM_) clusters. Analysis of CD8 T_CM_ clusters revealed significant differential gene expression patterns between the brain and spleen (**Figures 1H, S3E**). Notably, T_CM_8 C1 cells in brain tissues exhibited elevated expression of genes linked to T cell memory, including IL7R, the AP-1 transcription factors JUN and FOSB(39), and TCR signaling protein THEMIS associated with antigen-experienced T cell survival (40). Additionally, anti-inflammatory modulators ATF3, ZFP36L2, and NR4A2 were enriched in this brain cluster (41). Furthermore, genes such as GLUT3 (SLC2A3) regulating mitochondrial fitness and survival (42), and BTG1 linked to memory cell survival and quiescence were also more abundant in the brain T_CM_8 C1 cluster (43). Examination of marker genes within T_CM_8 C4 cells revealed heightened expression of cytolytic/resident markers GAMM, GZMK, CRTAM, and the inflammatory transcription factor NFκβ along with relatively lower IL-7R compared to C1 and C6 cells. Moreover, brain T_CM_8 C4 cells expressed genes induced upon TCR stimulation (SLC2A3 and NR4A2) at a higher level than their spleen counterparts, suggesting potential reactivation. Among T_CM_8 C6, the TNFα induced protein 3 (TNFAIP3) which inhibits IFNγ and TNFα in CD8 T cells, was significantly enriched in the brain relative to the spleen (44). Together, the brain CD8 T_CM_ clusters exhibit unique gene signatures associated with T cell memory, effector, and regulatory function.

Within the CD4 T cell lineages, 2 clusters were identified as CD4 T_CM_ subsets (C7 and C8). T_CM_ 4 C7 cells exhibited enrichment for the co-stimulatory molecules (CD28 and ICOS), IL-7R, transcription factors BACH2 associated with T cell survival (45), FOXP1 contributing to cell quiescence (46), PELI1 - a ubiquitin ligase negatively regulating T cell activation (47), memory genes (LTB, MAF, NFATC1), and β1 integrin (Figures1I, S3F). Cells in T_CM_4 C8, in addition to sharing expression of memory genes with C7 (IL7R, BACH2, LTB), expressed T_h_17 related genes (CCR6, AHR, RORA). Gene set enrichment analysis revealed concordance of gene signatures with longevity related, and MAPK pathways, while effector pathways such as NFκβ, RIG-I and TNF signaling were negatively represented (**Figure 1J-K**). In summary, our comprehensive scRNA-seq analysis of CD45+ cells unveiled a diverse range of T cell states including effector, central memory, and resident memory subsets in both the brain and spleen.

### T_CM_/T_RM_ loci are accessible in T cells within the brain

Given extensive research on CD8 T_RMs_ in the human and mouse brain (48–50), the identification of T cell subsets exhibiting T_CM_-like features sparked our interest. To explore the mechanisms regulating T_CM_ and T_RM_ differentiation states in the brain and validate our scRNA-seq data, we profiled the transcriptome and epigenome in parallel. We isolated nuclei from CD45+ and CD45- cells extracted from the brain (**Figure 2A**) and generated over 1.5 billion reads across 47,000 nuclei, with an average of 1378 genes/nuclei (**Figures S4A-D**). Transcriptome classification revealed a distinct network of cell clusters, including glial cells (microglia, oligodendrocytes), neurons, endothelial cells, cancer cells, and T cells (**Figures 2B, S4E-F**). The largest immune cluster comprised of macrophages, microglia, and T cells, with each cluster expressing genes encoding proteins with known cell-type distinctions. Specifically, cells in the macrophage cluster expressed CD86, TSPAN14, DOCK4, and TNFRSF21. The microglia cluster expressed ST6GALNAC3, ENTPD1, and P2RY12, while the T cell clusters were enriched for genes encoding (T_h_1 transcription factor [STAT4], T cell adaptor protein [SKAP1], as well as kinases and signaling molecules [TNIK, ITK, FYN]) (**Figures 2C,S4F**).

**Figure 2.**
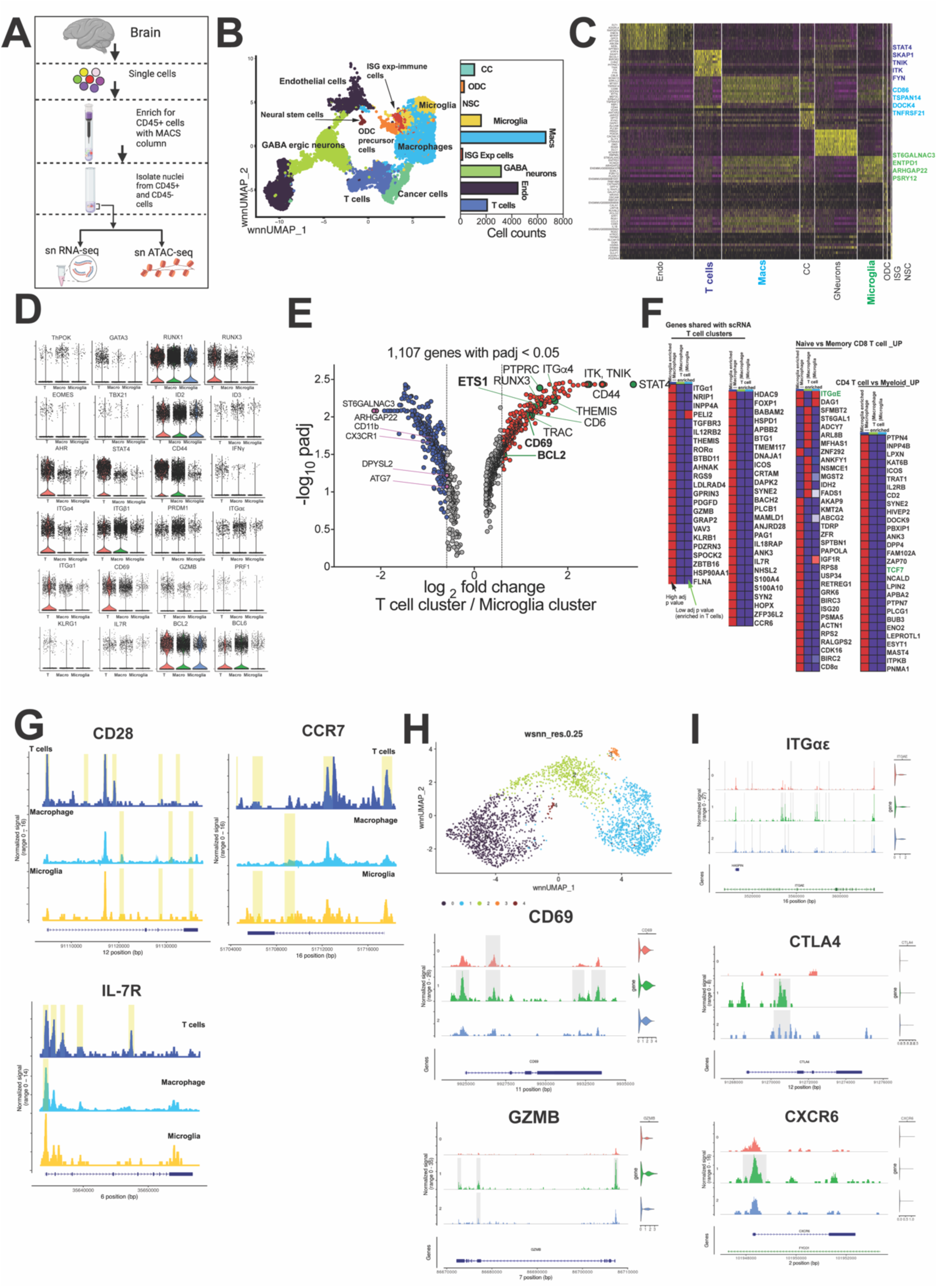
T_CM_/T_RM_ loci are accessible in T cells within the brain. **(A**) Schematic of analysis. **(B**) UMAP projection of 25,321 snRNA-seq profiles. Dots represent individual cells, and colors indicate cluster identity (labelled on right). **(C)** Heat map representation of RNA-seq of cluster-specific marker genes across all clusters. **(D)** Violin plots show expression of key genes across immune clusters. **(E)** Gene expression differences between T cell and microglial cell clusters. **(F)** GSEA of shared genes across sn and sc analysis**. (G)** Genomic regions showing snATAC-seq tracks of chromatin accessibility of T_CM_ genes across T cell, microglia, and macrophage immune clusters. **(H)** UMAP projection of T cell subclusters (2,158 T cells). **(I)** Genomic regions showing snATAC-seq tracks of chromatin accessibility of T_RM/EM_ genes across T cell clusters in H.

We identified the expression of several key genes that regulate T cell differentiation and function, including zinc finger transcription factors (TF) that regulate CD4 programs (ThPOK and GATA3), Runx TF (RUNX1 and RUNX3), T-box TF (EOMES and TBX21), inhibitor of DNA binding proteins (ID2 and ID3), TF regulating cytokine production (AHR and STAT4), markers of antigen-experienced cells (CD44, IFNG, ITGα4, and ITGβ1), markers of T cell residency (PRDM1, ITGα1, ITGαE, CD69, GZMB, KLRG1, and PRF1), and markers of long-lived cells with central memory features (BCL2 and BCL6) (**Figure 2D**).

To quantitatively evaluate genes enriched in T cells, we conducted differential gene expression analysis between T cells and microglial clusters. Within microglia, we discovered enrichment of canonical brain resident microglia transcripts, including ARHGAP22, DPYSL2, ATG7, CX3CR1, integrin ITAM (CD11b), transmembrane TMEM, and SIGLEC (51–53) (**Figure 2E**). In contrast, the genes that were highly expressed by the T cell cluster included those that regulate T cell signaling (TRAC, ITK, THEMIS, TNIK), transcription factors that control CD4 and CD8 T cell programs (STAT4, RUNX3), T cell migration (CD44, ITGα4), residency (CD69), and T cell survival (BCLA11B, BCL2), including the transcription factor ETS-1, which regulates the expression of IL- 7R. (54) (**Figures 2E, S4G**). Genes involved in CD4 T_h_17 function, RORα and AHR were also expressed in keeping with the transcriptome of T_CM_4 C8. Furthermore, a similar T cell gene expression profile was seen when comparing macrophages to T cells, with the IL-12-induced CD4 T_h_1 transcriptional regulator STAT4 being the most highly expressed gene in a significant fraction of T cells (**Figure S4H**). Using ATAC-seq analysis to identify regions of open chromatin, we found strong associations between chromatin accessibility of cis-regulatory elements within STAT4 and gene expression across clusters. Specifically, cells within the T cell cluster showed a high degree of chromatin accessibility, which corresponded with robust STAT4 expression (**Figure S4I**). Additionally, downstream targets of STAT4, including IFNγ and ICOS, showed distinct expression patterns in the T cell cluster compared to innate immune cells (**Figure S4J**). Interestingly, these genes were also among the top marker genes in the T cell cluster from our scRNA sequencing analysis of CD45+ cells.

To formally ascertain whether T cell clusters expressed genes that overlapped with our scRNA seq transcriptional profiles, we examined 7798 transcripts expressed by all three snRNA seq-derived immune clusters. We then used DEG p values for each expressed gene in T cells relative to macrophages and microglia. For comparison, we also included DEG p values of microglial transcripts relative to macrophages. Our gene set enrichment analysis revealed overlap with the top 20 marker genes expressed by T cell clusters from our scRNA seq, including classical effector/memory transcripts such as GZMB, CRTAM, and IL7R, among others (**Figure 2F**). Furthermore, when comparing the DEG genes to T cell signatures reported in murine models, we identified that ITGαE, the canonical integrin receptor of T_RM_, was enriched in T cells relative to macrophages (adjusted p value < 0.05). Additionally, we found that the transcription factor regulating central memory formation (TCF7) was highly enriched in T cells relative to microglia (adjusted p value = 1.76E-31) and macrophages (adjusted p value = 7.03E-13).

To assess T_CM_ gene accessibility, we focused on genes associated with T_CM_ function and longevity - CD28, IL7R, and BCL-2 - which were also expressed by macrophages and microglia. We found that there are more higher accessibility peaks in ATAC signals for CD28 and IL7R in T cells, indicating a more open chromatin state (**Figure 2G**), while BCL-2 accessibility was comparable across immune clusters (S4K). In particular, the higher accessibility peaks in CD28 and IL7R include a *cis*-regulartory element, the promoter region. Although CCR7 expression was relatively low, accessibility to CCR7 was significantly higher and distinct in T cell clusters compared to innate immune cells, where it was mostly absent (as in microglia). Furthermore, gene expression profiles and motif enrichment analyses using the hypergeometric optimization of motif enrichment (HOMER) database demonstrated that TFs regulating T_CM_ genes of the bZIP, RUNT, and ATF families were highly enriched and present in approximately 30% of target sequences within the T cell cluster.

To delineate genes regulating memory T cell states more precisely, we conducted further clustering analysis on 2,158 T cells, which resulted in the identification of four distinct clusters (**Figure 2H**). Since CCR7 expression was found in less than 1% of cells across all clusters, we focused our attention on T_RM_ genes and aimed to investigate whether the accessibility of promoter regions within genes controlling T_RM_ states was differentially regulated across the three major T cell clusters. Based on the proportion of cells expressing CD69 and GZMB, genes exemplifying T_RMs_, in Cluster 1 we hypothesized that Cluster 1 would show ATAC peaks of canonical T_RM_ genes. Correspondingly, visualization of peak tracks demonstrated greater chromatin accessibility within gene regulatory regions of CD69, GZMB, ITGαe, CTLA4, and CXCR6 in Cluster 1 (**Figure 2I**). Motif analysis showed close to 25% of sequences within these T clusters contained motifs for Blimp1 (encoded by PRDM1), canonical transcriptional repressor of residency fate in memory T cells (p = 1.0 E-5).

Overall, the data revealed that T cells display distinct chromatin accessibility patterns for genes regulating T cell residency and lymph node homing ability indicating that T cells with putative resident and central memory features populate the primate brain.

### CCR7^+^ CD4 T cells in CNS share phenotypic features with T_CM_ in blood and lymph nodes

Single-cell gene expression computational analyses revealed two distinct T cell subsets in the healthy brain: a tissue-resident subset expressing CD69, and a central memory subset expressing IL-7R and BCL-2, with accessible CCR7 locus. To validate and extend our findings from single cell sequencing, we investigated the immune makeup of the leptomeningeal space by analyzing cerebrospinal fluid (CSF). CSF samples were collected from the foramen magnum, alongside paired axillary lymph node aspirates and blood samples (**Figure S5A**). Unlike blood, the CSF exhibited minimal presence of B cells and monocytes, indicating preferential infiltration of T cells into the CSF under steady-state conditions.

CD28 and CD95 expression analysis in T cells revealed enrichment of CD28^high^ CD4 memory T cells in the CSF, while terminally differentiated (CD28- CD95+) and naive (CD28^int^ CD95-) subsets were notably absent, consistent with reported human phenotypes (55). Among CD8 T cells, the CD28^high^ CD95+ population was similar, but CD28- CD8 T cells constituted approximately 18% of the CSF CD8 T cell pool on average (range: 10-25%, **Figure S5B**). We validated detection of CCR7 expression at 4°C, considering the standard practice of staining at 37°C in mice cells (data not shown). Using the CD28^high^ subset in blood as a reference, our analysis revealed approximately 50% co-expression of CCR7 in CD28^high^ CD4 T cells within the CSF, while CCR7 expression in CD28^high^ CD8 T cells within the CSF averaged around 20% (**Figure S5C**).

To gain detailed understanding of T cell phenotypes across distinct anatomic sites, we compared phenotype of CSF-derived cells to those in matched CNS tissues and adjacent lymph nodes, spleen, and blood. T cells in the brain exhibited distinct differentiation states; while CD28^high^ CD4 T cells were enriched in the CNS, in keeping with their transcriptional profiles, CD8 T cells were predominantly CD28- (**Figure 3A**). Similar to the CSF, CD28^high^ CD4 T cells in the parenchyma exhibited heterogeneity in terms of CCR7 expression (**Figures 3B**). We compared CCR7+ CD4 T cells in the CNS (choroid plexus and brain parenchyma) to those in corresponding lymphoid compartments (deep cervical lymph nodes and spleen) to explore potential phenotypic similarities to T_CM_. Analysis of receptor expression revealed reduced per cell CCR7 levels in the CNS compared to lymphoid counterparts. (**Figure 3C**).

**Figure 3.**
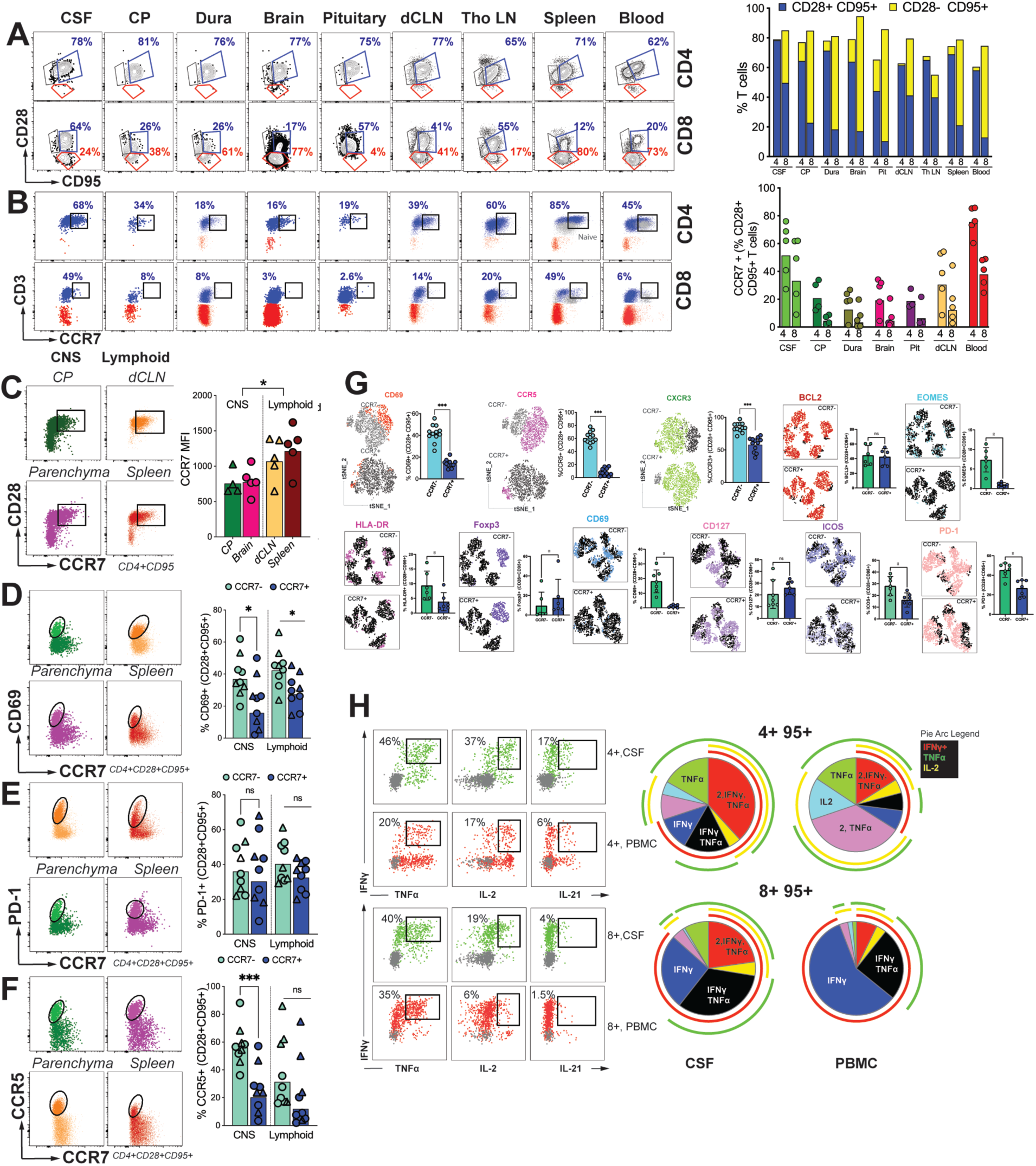
CCR7^+^ CD4 T cells in CNS share phenotypic features with T_CM_ in blood and lymph nodes. **(A)** Representative flow plots identifying CD28 and CD95 expression on CD4 (top row) and CD8 (bottom row) T cells from the CSF, CP, Dura, Brain, Pit, dCLN, Th LN, Spleen, Blood (Left); Frequencies of CD28+CD95+ (blue) and CD28-CD95+ (yellow) in CD4 T cells and CD8 T cells (right). **(B)** Representative flow plots indicating CCR7 expression on CD28^High^ CD4 (top row) and CD28^High^ CD8 (bottom row) T cells (left); frequencies of CCR7 expression on CD28^High^ CD4 and CD28^High^ CD8 T cells from the CSF, CP, Dura, Brain, Pit, dCLN, Th LN, Spleen, Blood. **(C)** Representative flow plots indicating CD28 expression and CCR7 expression on CD4+CD95+ T cells in the CNS (Parenchyma and CP) and the Lymphoid tissues (dcLN and Spleen) (Left); CCR7 MFI of CD4+CD95+ (Right). **(D**) Representative flow plots indicating CD69 expression and CCR7 expression on CD4+CD28+CD95+ T cells in the CNS (Parenchyma and CP) and the Lymphoid tissues (dcLN and Spleen) (Left); frequency of CD69+ on CD4+CD28+CD95+ CCR7-/+ T cells (Right). **(E)** Representative flow plots indicating PD-1 expression and CCR7 expression on CD4+CD28+CD95+ T cells in the CNS (Parenchyma and CP) and the Lymphoid tissues (dcLN and Spleen) (Left); frequency of PD-1+ on CD4+CD28+CD95+ CCR7-/+ T cells (Right). (**F**) Representative flow plots indicating CCR5 expression and CCR7 expression on CD4+CD28+CD95+ T cells in the CNS (Parenchyma and CP) and the Lymphoid tissues (dcLN and Spleen) (Left); frequency of CCR5+ on CD4+CD28+CD95+ CCR7-/+ T cells (Right). **(G)** Representative tSNE plot illustrating CD69, BCL2, CD69 (column 1), CCR5, EOMES, CD127 (Column 2), HLA-DR, ICOS (Column 3), and CXCR3, Foxp3, PD-1 (Column 4) expression on CD4+CD28+CD95+ CCR7-/+ T cells in the CSF, Blood and PBMCs; frequencies for each population are expressed to the right of the tSNE plots. **(H)** Representative flow plots illustrating cytokine (IFNγ, TNFα, and IL-2) production in the CSF and PBMCs after intracellular cytokine stimulation (ICS), pie charts on the right.

In each tissue, CCR7+ CD4 T cells displayed a notably lower likelihood of expressing the typical T_RM_ marker, CD69 (**Figure 3D**). The presence of the inhibitory receptor PD-1, observed in CD8 T_RM_ (56), showed similar levels between subsets (**Figure 3E**), implying comparable local TCR stimulation. Given reports of CCR5 expression in CD69^+^ CD4 T_RM_’s in the intestinal mucosa (57), we investigated if CCR7- CD4 T cells, with higher CD69 expression, exhibited proportionally elevated CCR5 expression. Our findings confirmed this phenomenon within the CNS, but not in lymphoid tissues (**Figure 3F**). In essence, the data indicate that CD4 T cells within the non-inflamed brain parenchyma exhibit CD28^high^ expression and diverge based on CD69 and CCR7 levels, with CCR7+ T cells sharing traits with their lymphoid counterparts.

To delve into the phenotypic distinction between CCR7 subsets, we first studied T cells in the CSF. Flow SOM clustering on live CD4+ T cells (from n=4) was first performed to determine if CCR7+ and CCR7- subsets represent distinct clusters. Based on expression of specific T_h_1 (CXCR3, CCR5), T_h_17 (CCR6), activation and memory markers (CD95, PD-1, CD69), 5 metaclusters were defined with varying levels of CCR7 expression (**Figure S6A**). CCR7- and CCR7+ clusters showed enrichment of distinct surface markers; CCR5 and PD-1 were enriched in the CCR7- cluster, while CCR6 was enriched in the CCR7+ cluster (**Figure S6B**).

Subsequently, we investigated whether CCR7+ CD4 T cells in the CSF displayed features typical of quiescent T_CM_, and conversely, whether CCR7- CD4 T cells exhibited characteristics akin to T_RM_ or activated effector memory cells. Notably, while CD4 T cells in blood lacked CD69 expression, CD69 was predominantly expressed within the CCR7- subset of CSF CD4 T cells, mirroring the expression pattern observed within the brain parenchyma (**Figure 3G**). Likewise, the expression of CCR5 and CXCR3 was primarily limited to the CSF CCR7- subset. This distribution was also consistent for markers associated with activation (ICOS, EOMES, PD-1, and HLA-DR), while Foxp3, a transcription factor for T regulatory cells, exhibited enrichment in CCR7+ cells. In further support of CCR7+ cells representing resting memory cells, we observed the presence of CD127 and BCL-2 expression, although at levels comparable to CCR7- populations (**Figure S7**).

In summary, our findings demonstrate that CCR7+ CD4 T cells in the CSF and brain parenchyma share phenotypic similarities with those in lymphoid tissues, showcasing core traits of T_CM_. Accordingly, we hypothesized that CNS CD4 T cells would display a functional hallmark of T_CM_ by producing IL-2. Stimulation of CSF CD4 T cells with PMA/Ionomycin revealed that CD4 T cells were polyfunctional and produced IL-2. Conversely, a smaller proportion of IL-2+ CD8 T cells was identified, consistent with the lower relative frequencies of CCR7+ CD8 T cells in CSF (**Figure 3H**). Altogether, our phenotypic data support the conclusion that CCR7^+^ CD4 T cells in the CNS share phenotypic features with traditional T_CM_ found in the periphery.

### Sequestration of CD4 T_CM_ in lymphoid tissues reduces CCR7^+^ CD4 T cell frequencies in CSF and elevates soluble inflammatory markers

We next investigated whether CNS CCR7+ CD4 T cells displayed typical migration patterns to and from lymphoid tissue, a hallmark of T_CM_ cells. To test the hypothesis that sequestering T_CM_ cells in lymph nodes would decrease frequencies of CCR7+ CD4 T cells in CSF, we dosed rhesus macaques (n = 12) with FTY720 (30 μg/kg/day) for 4 weeks and analyzed paired blood and CSF T cells (**Figure 4A**) for up to eight weeks to capture T cell dynamics within the subarachnoid space (SAS) during FTY-induced peripheral lymphopenia and lymphocyte rebound following drug cessation. Analysis of blood T cells showed a rapid and dramatic decline in total CD4 T cells in the periphery within 1 week after treatment, while CD4 T cell counts significantly decreased in the CSF at week 4 post FTY720 treatment (**Figure 4B**). Consistent with FTY720 mediated inhibition of S1PR-mediated T cell egress and retention of CCR7-expressing T cells in lymph nodes, naive T cells, subsets with the highest per cell expression of CCR7, were rapidly sequestered as evident by a 4-fold decline in absolute counts of naive (CD28+ CD95-) T cells within 1 week of FTY720 treatment. By week 4, naive T cells declined further with a 100-fold drop in CD4 T cells and 80-fold decline in CD8 T cells relative to baseline levels (**Figure S8A**). In contrast, it took up to 2 weeks for the peripheral CD4 T_CM_ pool to decrease significantly and by week 4, a 17-fold decrease in CD4 T_CM_ was observed. This decline was accompanied by a significant decrease in CCR7+ CD4 T cells in CSF by week 4 indicating direct recruitment of CCR7+ CD4 T cells into the SAS from lymphoid tissues via the systemic compartment.

**Figure 4.**
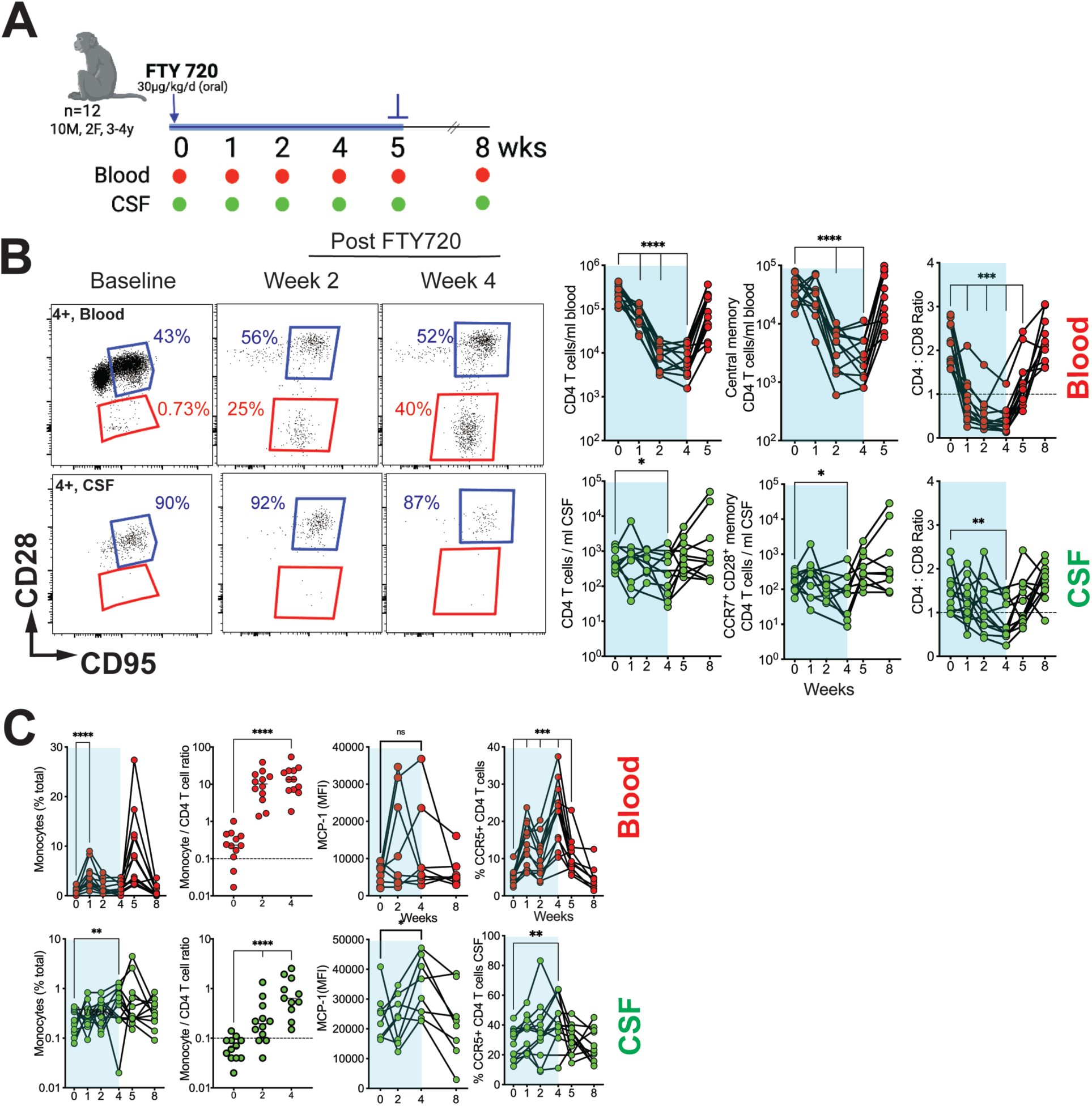
Sequestration of CD4 T_CM_ in lymphoid tissues reduces CCR7^+^ CD4 T cell frequencies in CSF and elevates soluble inflammatory markers. **(A)** Study schematic: n= 12 rhesus macaques between the ages of 3-4 years were administered an oral dose of 30 ug/kg per day Fingolimod (FTY) for the first four weeks of the study. CSF taps and blood draws were performed at indicated timepoints. **(B)** Representative longitudinal flow plots indicating CD28 and CD95 expression on CD4 T cells from the blood (top row) or the CSF (bottom row) (Left); CD4 T cell counts/ml, Central memory CD4 T cells and CCR7+CD28+ memory CD4 T cells/ ml blood or CSF, and CD4:CD8 Ratio for Blood and CSF (Right). **(C)** Frequencies of Monocytes, Monocyte to CD4 T cell ratio, Median Fluorescent Intensity (MFI) of Monocyte Chemoattractant Protein-1 (MCP-1), and CCR5 expression of CD4 T cells in the blood and CSF over the course of the study.

While T_CM_ CD8 T cells also showed a similar decline in the periphery (**Figure S8B**), total CD8 T cells within the CSF were not significantly diminished following FTY treatment. Indeed, the relative stability of CD8 T cells over CD4 T cells in the CSF led to a significant decrease in CD4:CD8 ratio in CSF at week 4. Strikingly, the frequency of CD28^-^ effector memory cells increased at week 1 both for CD4 and CD8 T cells in blood [CM:EM ratios decreased from 34 to 6 for CD4 T cells at week 1 and week 4, and EM:CM ratio increased from 2.2 to 10.4 for CD8 T cells]. Despite the dramatic shift in the CM:EM ratio of T cells in blood, the CSF T cell CM:EM ratio remained stable over the 4-week period, and we observed little to no increase in CD28- CD95+ CD4 T cells in the CSF demonstrating that composition of T cells is stringently regulated to exclude CD28- CD95+ CD4 T cells from the SAS.

Since studies in mice demonstrate that T cell elimination from the meninges can result in pro-inflammatory innate immune skewing (58), we assessed monocyte frequencies to determine if a similar compensatory increase in monocytes might be observed within the SAS following low dose FTY720 treatment. The data showed a net increase in monocytes in the CSF at week 4, resulting in a significant increase in monocyte/CD4 T cell ratio in the CSF (**Figure 4C**). Significant elevation of monocyte chemotactic protein-1 (MCP-1) at week 4 in the CSF but not in the blood suggested CSF monocyte influx was chemokine mediated. We also noted increased levels of interferon protein 10 (IP-10), but pro-inflammatory cytokines/chemokines IL-6, IL-8, and IL-1β remained unaltered in the CSF arguing against a generalized pro-inflammatory state during a window of CCR7+ CD4 T cell reduction in CSF (data not shown). In conclusion, the data strongly suggest that CCR7+ CD69- CD4 T cells in the CSF are predominantly recruited from lymphoid tissues. Moreover, the identification of immune activation associated with a reduced presence of CCR7+ CD4 T cells in the SAS indicates a potential role for these cells in maintaining neuroimmune homeostasis.

### CCR7^+^ CD4 T cells in CNS exhibit functional T_CM_ features and reside within skull bone marrow

Given the significance of bone marrow for T_CM_ localization, particularly the skull’s hematopoietic-rich marrow cells (59, 60), we investigated whether CD4 T cells with T_CM_ attributes reside there. After manually extracting marrow cells from the skull, single-cell suspensions were stained to identify innate and adaptive immune cells. Consistent with the distribution observed in the mouse brain, analysis revealed three distinct subsets among CD3- CD45+ cells based on their expression of CD11b and HLA-DR markers (**Figure S9**). Apart from myeloid cells, the immune environment within the skull bone marrow also included T cells, constituting 71% of CD45+ cells, which is similar to the frequencies observed in the CSF (**Figure 5A**). Furthermore, the skull bone marrow exhibited a higher proportion of CD8 T cells relative to CD4 T cells, akin to the composition in the brain parenchyma (**Figure 5B**). With respect to memory cells, similar to the CNS, the expression of CCR7 was heterogeneous within the CD28+ CD4 T cell population, whereas CD28- CD4 T cells predominantly lacked CCR7 expression (**Figures 5C-D, S10**).

**Figure 5.**
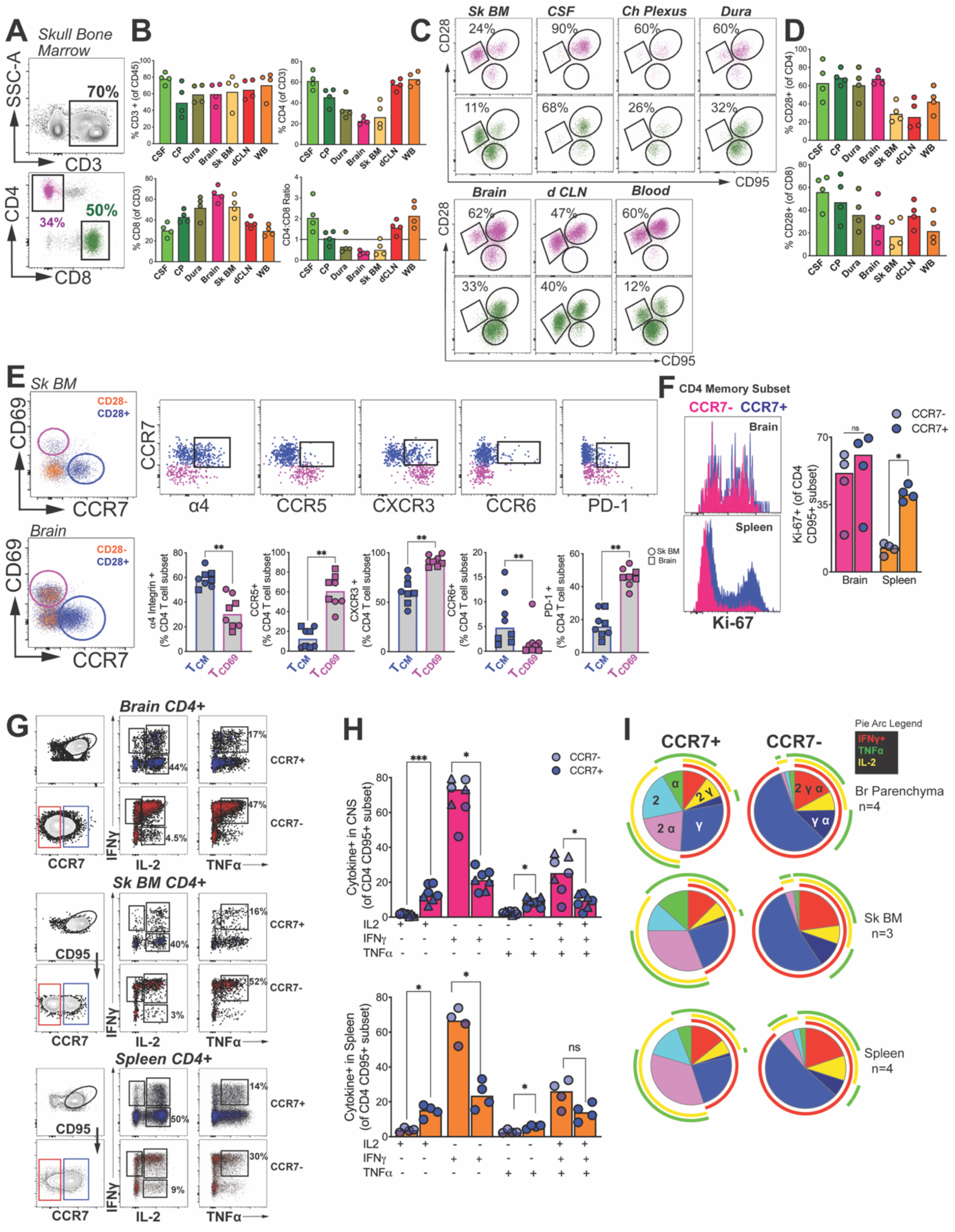
CCR7+ CD4 T cells in CNS exhibit functional T_CM_ features and reside within skull bone marrow. **(A)** Gating within the skull bone marrow and **(B)** corresponding frequencies of CD3, CD4 (top), CD8 T cells, and CD4/CD8 ratios (bottom) across tissue compartments (Cerebrospinal fluid [CSF], Choroid Plexus [CP], Dura Mater [Dura], Brain, Skull Bone Marrow [Sk BM], drain cervical lymph node [dCLN], and whole blood [WB]). **(C)** Population gates for CD4 (purple) and CD8 (green) subsets (naive [CD28+CD95-], central memory [CD28+CD95+], effector memory [CD28-CD95+]) with **(D)** corresponding frequencies of CD28+ subsets across tissue compartments. **(E)** Median fluorescent intensity of CD69 (top) and CCR7 (bottom) expression within CD4 T cell subsets across tissue compartments. **(F)** Phenotypic characterization (integrins: a4; chemokine receptors: CCR5, CXCR3, CCR6; activation markers: PD-1) of Tcm-like (CCR7+) and tissue resident (CD69+) CD4 T-cells from brain and skull bone marrow. **(A-F)** Data points indicate individual tissue samples. **(F)** Symbols indicate skull bone marrow (circle) or brain tissue (square) derived samples. Bars indicate medians.

Additional phenotypic analysis based on CCR7 and CD69 expression demonstrated that the T_CM_ subset exhibited distinct phenotypes compared to the T_RM_ subset in both the skull bone marrow and brain. Specifically, the T_CM_ subset had higher relative expression of integrin α4 and lower expression of chemokine receptors CCR5 and CXCR3. Moreover, T_CM_ cells displayed elevated CCR6 expression, while PD-1 expression was more prominent in T_RM_ cells (**Figure 5E**). Functional characterization of memory CCR7- and CCR7+ CD4 subsets demonstrated that both brain CCR7- and CCR7+ CD4 T cells were able to equally mount recall proliferation ex vivo, while splenic CCR7+ CD4 T cells, and to a lesser extent CCR7- CD4 T cells were Ki67+ (**Figure 5F**). Analysis of cytokine production following PMA/I stimulation showed that CNS CCR7+ CD4 T cells produced a higher relative frequency of IL-2, while the CCR7- subset produced high levels of IL- 2, IFNγ and TNFα. This cytokine expression pattern was consistent with T_CM_ functionality, as exemplified by a similar cytokine pattern in splenic CD4 T_CM_ cells (**Figure 5 G-I**). In summary, our findings highlight the presence of a CCR7+ CD4 population in the brain and skull bone marrow that exhibits T_CM_-like characteristics, akin to T_CM_ in the spleen.

### vRNA within the brain encompassing frontal and temporal lobes during chronic SIV infection

To better understand the role of CD4 T_CM_ in the CNS, we investigated these cells in a well-established model of chronic viral neuroinflammation (61). Aged rhesus macaques (17-20 years old) were infected with the neuropathogenic SIV strain (SIVCL757) to replicate key aspects of chronic neuroinflammation and establish CNS virus presence. Following the post-acute phase, to ensure CNS viral dissemination, intermittent antiretroviral therapy (ART) was initiated (weeks 16-52 post SIV when vRNA in CSF and plasma were both above threshold of detection, i.e., > 15 vRNA copies/ml). We adopted this treatment regimen to induce cycles of viral suppression and rebound within the CNS, simulating scenarios in individuals at risk of neurological co-morbidities due to chronic neuroinflammation [23, 24] (**Figure 6A**). Longitudinal collections of CSF and matched blood was conducted from infected animals for up to 116 weeks, except for one SIV+ animal (34974) that was euthanized at 52 weeks due to health complications. Prior to necropsy, ART was interrupted in all animals except 33191, which was ART naive, to provoke viral rebound. At necropsy, CNS and peripheral lymphoid tissues were collected for analysis, assessing viral loads, lymphocyte subpopulation alterations via flow cytometry, single-cell transcriptomics of brain leukocytes, and spatial transcriptomics of the hippocampus. An age-matched control group of SIV-unexposed animals (n=5) was also studied.

**Figure 6.**
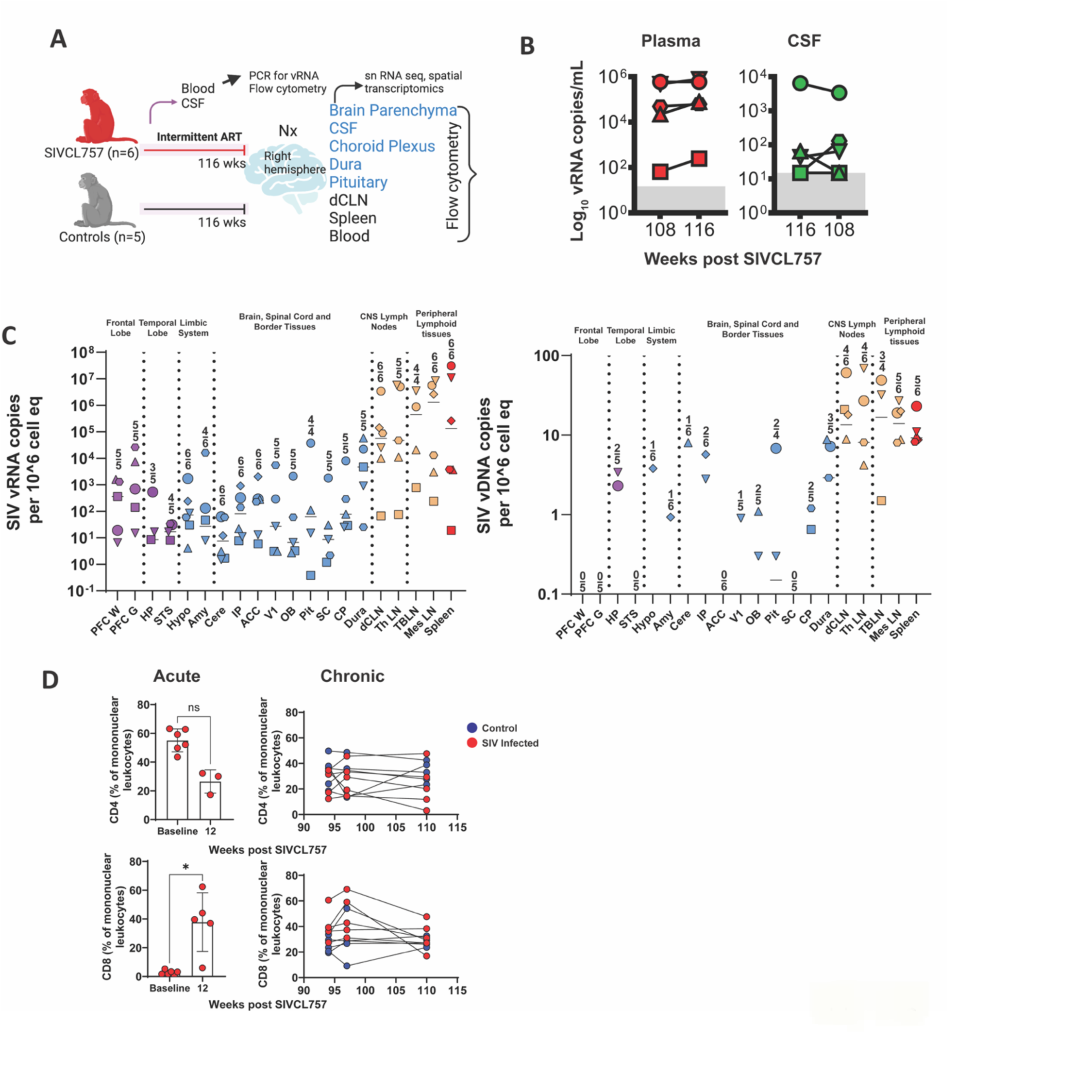
vRNA within the brain encompassing frontal and temporal lobes during chronic SIV infection. **(A)** Study schematic: rhesus macaques were either infected with SIVCL757 intravenously or age matched uninfected controls were longitudinally assessed over the course of the study to assess longitudinal peripheral and CNS viral burden. At necropsy, various CNS tissues (Brain parenchyma, CSF, Choroid Plexus, Dura, Pituitary, dCLN, Spleen and Blood) were assessed to measure tissue viral burden. Single cells isolated from the brain were utilized for single nuclei (sn) RNA sequencing. Brain tissue from the Hippocampal region was utilized for spatial transcriptomics. **(B)** Kinetics of plasma (red) and CSF (green) viral loads during the chronic phase (wk108 – 116) of SIVCL757 infection. **(C)** vRNA and vDNA in various brain regions, dura mater, deep cervical lymph nodes, and PBMCs. **(D)** CSF CD4 and CD8 frequencies during the acute phase (12 weeks post infection) and chronic phase (week 92-110) of SIVCL757 infection.

Following infection, viremic animals (n=4) exhibited median plasma viral loads of 165,000 copies/mL at week 3, with CSF viral RNA (vRNA) reaching a median of 19,750 vRNA copies/mL (**Figure S11**). vRNA in both plasma and CSF exhibited a lower magnitude and variable pattern when compared to viral loads observed following SIVmac251 infection (62). Similar to SIVmac251 infection, there was plasma-CSF concordance during the acute SIVCL757 infection before initiation of ART. However, there was an exception in one TRIM5a-restrictive animal (32967), which displayed transient plasma viral discordance up to week 6 post-infection. Of note 2 animals (33191 and 34996) demonstrated sporadic and minimal vRNA in CSF despite plasma vRNA after the acute phase. ART initiation between sixteen to forty-six weeks post-infection led to viral suppression (vRNA copies <15) in plasma as early as 4 weeks and as late as 6 weeks post ART. Throughout chronic infection, viral loads in CSF were consistently 3 log-fold lower than those in plasma (median viral loads/ml at week 108: plasma, 50,000; CSF, 65), aligning with our previous findings in acute SHIV.C.CH505 infection (**Figure 6B**) [25].

At necropsy, 3mm post-mortem punch biopsies were collected from specific brain regions to assess vRNA and vDNA in various brain regions, border tissues, CNS-draining lymph nodes, and peripheral lymphoid tissues (**Figure 6C**). The frontal lobe, linked to cognition, displayed vRNA positivity in both gray and white matter regions across all animals tested. However, vDNA was not detectable, falling below the detection limit. The detection of vRNA and vDNA exhibited variability in the temporal lobe, limbic system, and other brain and border tissues. While the CNS-draining lymph nodes and peripheral lymphoid tissues showed vRNA in all animals, vDNA was not consistently detected across these tissues in certain animals. These observations imply ongoing viral replication within the CNS, with limited evidence of viral integration in assessed CNS tissues.

The presence of virus within the rhesus brain prompted investigation into acute or chronic alterations in CSF CD4 or CD8 T cell populations. Lymphocyte analysis of the CSF showed a trend of CD4 T cell reduction during the initial 12 weeks of infection (not significant), while CD8 T cells exhibited an increase (p<0.05; fold change: 13). Both CD4 and CD8 T cell frequencies stabilized during the chronic phase of infection (**Figure 6D**). These findings highlight widespread vRNA in the CNS, low vDNA levels, and acute changes in CD4 and CD8 T lymphocyte populations within the CSF following SIVCL757 infection.

### Spatial profiling of hippocampus shows induction neuroinflammatory and neurodegenerative gene programs during chronic SIV infection

The presence of vRNA in the chronically infected macaque brain, prompted us to investigate inflammatory pathways using spatial transcriptomics. We initiated our investigation by examining T cell distribution in the human brain. Through immunohistochemistry (IHC) analysis of paraffin-embedded brain hippocampal sections from both glioblastoma patients (GBM-01) and non-demented individuals (90-128, 06- 080, 98-0189, and 06-080) sourced from the Netherlands Brain Bank, we aimed to elucidate the presence and localization of neurons (NeuN), myeloid cells (CD11b, IBA1), and lymphocytes (CD45, CD3, CD4). Tonsil sections from a healthy individual exhibited abundant T cells and myeloid cells, without neuron-specific staining. In contrast, hippocampal sections from non-demented patients displayed microglia, neurons, T cells, and monocytes, primarily around blood vessels (**Figure S12**). Comparatively, glioblastoma patient-derived hippocampal tissue sections exhibited pronounced T cell distribution throughout the brain parenchyma.

Having established the presence of T cells in the human brain with IHC, our focus shifted to assessing the histological and T cell landscape within the hippocampal tissue of chronically SIV- infected macaques. This investigation was carried out on hippocampus tissue samples from two animals – one healthy control (33980) and one SIV+ animal (35595) using the Nanostring Digital Spatial Profiler (DSP) platform. We employed CD3, CD45, and NeuN as morphological markers to identify T cells, leukocytes, and neurons. By evaluating the expression of CD45, CD3, and NeuN, we selected 24 regions of interest (ROIs) that encompassed varying levels of surface CD45 expression, covering distinct spatial zones within the hippocampus. These zones comprised regions adjacent to Cornu Ammonis, as well as areas around small and large blood vessels and parenchymal regions (11 ROIs and 10 ROIs respectively) (**Figure 7A**). Subsequently, these 24 ROIs underwent 147-plex antibody profiling and whole transcriptomic analysis (WTA) for comprehensive characterization.

**Figure 7.**
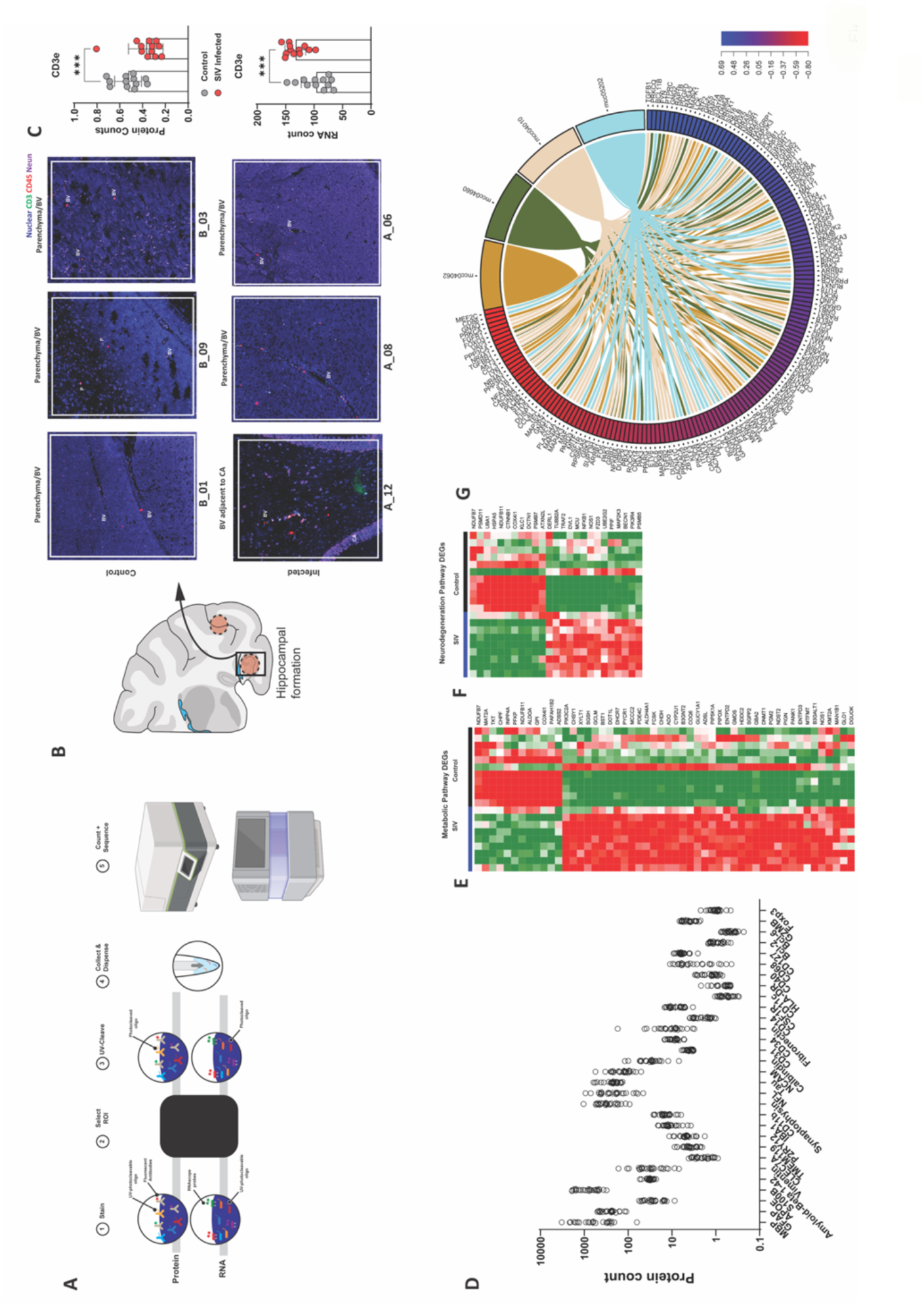
Spatial profiling of hippocampus shows induction neuroinflammatory and neurodegenerative gene programs during chronic SIV infection. **(A)** Study schematic for Nanostring whole transcriptome analysis (WTA) and proteomics pipeline. **(B)** Representative illustration for ROI selection within the hippocampal region of control (top) and SIVCL757 infected (bottom) animals; Nuclear (blue), CD3 (green), CD45 (red), and NeuN (Purple) **(C)** CD3ε Protein and RNA counts for ROIs. **(D)** Multiple Protein counts for ROIs. **(E)** Differentially expressed metabolic genes between control and SIV infected ROIs. **(F)** Differentially expressed neurodegenerative genes between control and SIV infected ROIs. **(G)** Chord plot for differentially expressed genes between control and SIV infected CD45-enriched cells from single cell transcriptomics.

The expression CD45, CD3, and NeuN proteins displayed heterogeneity across the 24 ROIs selected. By utilizing fluorescence signal of CD3 and CD45, we identified T cells primarily within blood vessels. Notably, both CD3 and CD45 protein signal were above the limit of detection in our analysis (**Figure 7B and 7C**). Overall, we observed lower CD3ε protein counts, but higher RNA counts in SIV infected ROIs. Our antibody profiling demonstrated expected enrichment of protein signals corresponding to glial cells (oligodendrocytes [myelin basic protein], astrocytes [GFAP, APOE, S100B, amyloid β, Vimentin], microglia [CLEC7A, TMEM119, CD11b, IBA1, P2RY12]) neuronal proteins (synaptophysin, neurofilament light chain, Tau, NCAM [CD56], Vimentin, Calbindin). Proteins expressed by endothelial and muscle cells [CD31, CD34, Fibronectin] were also detected. With respect to immune proteins, signal for myeloid cells (CD14, CSF1R, CD11c, HLA-DR, CD40, CD68) was observed. In keeping with the transcriptomic data, expression of T cell identity proteins was observed - memory T_CM_ cells (CD127, Bcl-2, Bcl-6) effector/resident cells (GZMB) and transcriptional regulators (BCL-6, FOXP3) (**Figure 7D**).

### SIV infection induces metabolic shifts within the hippocampus

Differential expression analysis between control and SIV-infected ROIs (n =12 control; n=12 SIV) revealed altered expression of metabolic and neurodegenerative genes in response to SIV. During neuroinflammation, microglia shift their metabolism from oxidative glycolysis to aerobic glycolysis, essential for cytokine release (63). Given the enrichment of microglial proteins in our protein analysis, we investigated metabolic shifts in our RNA analysis of 24 ROIs, and found 12 downregulated and 40 upregulated metabolic genes in response to chronic SIV (**Figure 7E**). Notably, numerous genes regulating methionine-related processes were altered; we identified downregulation of MAT2A and upregulation of ALDH4A1 and CHDH. In vitro studies have shown that MAT2A inhibition promotes neurological recovery(64), and changes in choline dehydrogenase (CHDH) are observed in the AD hippocampus (65). Neuroinflammation also impacts mitochondrial electron transport chain (ETC) genes in conditions like traumatic brain injury (66), Alzheimer’s disease (AD) and Parkinson’s disease (PD) (67, 68). Specifically, we saw decreased expression of ETC genes, NDUFB7, NDUFB11, and COX4I1, along with a decrease in inositol polyphosphate phosphatase 4A (INPP4A), a suppressor of glutamate excitotoxicity in the CNS linked to neurodegeneration in the striatum (69). We identified an increase in BST1, a risk factor for neurodegenerative diseases, particularly PD (70–75). Additionally, there was an increase in Dot1L, which regulates receptor-interacting protein kinase 1 (RIPK1) and triggers apoptotic death during cerebral ischemia injury. PDE4C, associated with regulating inflammatory cell activation and commonly expressed in myeloid lineages within the brain, showed increased expression (76). Methionyl-tRNA formyl transferase (MTFMT) instrumental in regulating mitochondrial function to mediate innate defense against infection was increased (77). Nitric oxide synthase (NOS1), associated with AD, psychiatric disorders, and behavioral deficits (78) also displayed increased expression. An increase in Lysine methyltransferase 2A (KMT2A), which plays a critical role in monocyte and macrophage inflammation, was also noted (79). Thus, with chronic SIV infection, significant metabolic shifts, including altered expression of genes related to methionine processes, mitochondrial electron transport chain, and neurodegenerative risk factors, occur highlighting intricate molecular changes associated with neuroinflammation.

### Activation of neurodegenerative and neuroinflammatory gene programs in SIV infected brain

We examined differential gene expression associated with neurodegenerative processes in SIV-infected brain regions and observed downregulation of HSP5, CTNNB1, COX4I1, KLC1, DCTN1, and PSMB7. These genes are linked to various neurodevelopmental and neurodegenerative disorders including AD and PD (80). For instance, COX4I1 is downregulated in entorhinal and hippocampal tissues of AD patients, KLC1 is downregulated in AD (81), DCTN1 deficiency is connected to PD (82), and PSMB7 downregulation is implicated in AD (83). Furthermore, we observed an increase in TRAF2, NFκB1, FZD3, MAP2K3, and autophagy- related genes BECN1 and PIK3C3, which have known impacts on immune response, inflammation, and neurodegeneration. TRAF2 is involved in apoptotic suppression and necroptotic signaling by TNFα while also negatively regulating cerebral ischemic injury (84, 85). The upregulation of the mitochondrial calcium uniporter, located on the inner mitochondrial membrane, is noteworthy, as disturbances in calcium homeostasis are linked to AD and PD (86–90). The upregulation of FZD3 is significant; when FZD combines with Wnt proteins on microglia, it initiates the canonical Wnt pathway, leading to microglial activation (91–95). Additionally, the upregulation of MAP2K3, a dual specificity kinase activated by stress, is interesting as it is enriched within microglia in the mouse cortex, upregulated in AD populations, and its signaling can contribute to neuroinflammation(96–98). The increase in BECN1 and PIK3C3, essential components of the autophagy multi-protein complex, is noteworthy; these genes have been implicated as key risk factors for AD and have been shown to exacerbate neurodegeneration when proteolytically cleaved(99, 100).

Furthermore, the notable increase in the inflammatory mediator NFκB1 is of significance due to its role in promoting T_h_1 cell differentiation and inducing cytokine production in innate immune cells, such as IL-12, which drives T_h_1 fate (101, 102). Shifting our focus to immune-regulating genes within the Hippocampus during chronic neuroinflammation, we observed an increase in TIE1 (Log2 FC - 0.53), a tyrosine kinase that enhances adhesion of monocytes to endothelial cells via upregulation of cell adhesion molecules. TIE1 has also been noted to increase in the brains of HIV+ individuals with neurocognitive impairment. Similarly, the macrophage specific gene, MPEG1, upregulated in highly active MS and in response to proinflammatory signals such as TNFα and LPS, was also induced. Although a significant increase in TNFα within the CSF was not observed during chronic SIV, EIF4E (Log2 FC - 0.39), a eukaryotic initiation factor, was elevated; in the brain, EIF4E controls the translation of mRNAs related to the inflammatory response, including IκBα, a repressor of the NF-κB which regulates TNFα expression. Lastly, we found that the interferon-induced protein IFIT3 (Log2 FC - 1.05), induced upon viral infection, was also induced, aligning with our observations of vRNA and vDNA in the white and grey matter of the Hippocampus.

### Single cell analysis identifies activated inflammatory macrophage population in SIV infected brain

Expanding upon our hippocampal spatial transcriptomics data, we employed an additional analytical approach involving single-cell gene expression data of CD45+ enriched brain cells from controls (33980, 33994), described in Figure 2, alongside SIV+ animal (32967). Analysis of differential gene expression from this single-cell dataset unveiled notable alterations in pathways encompassing chemokine, T cell receptor, MAPK signaling, and transcriptional misregulation in cancer due to chronic SIV infection. Key genes linked to T cell function (STAT4, PTPN6, NFATC3, NF-κB1, JAK3, GRAP2, TGFβR2, CDK4, NF-κBIA), MAPK signaling pathway (EPHA2, PTPRR, TRAF2, NFATC3, NF-κB1, MAP2K3, TGFβR2, MAP3K4, MAP3K13, TAOK1, MAPK8IP3), and Transcriptional misregulation in cancer (SPI1, NFATC3, NF-κB1, DOT1L, TGFβR2) were enriched in Hippocampal ROIs of chronically infected animals (**Figure 7G**).

To delineate microglial or macrophage activation, we initiated subclustering of myeloid cells into eight distinct subclusters, each with distinct phenotypes. Utilizing well-established microglial markers (PTPRC, ITGAM, CX3CR1, P2RY12, and TMEM119), cluster 3 and cluster 5 emerged as microglia-like (Figure 8A-B), while clusters 0, 1, 2, 4, and 7 expressed definitive macrophage markers CD68 and FCGR1A. HLA genes B2M and CD74 were primarily expressed in clusters 0, 2, 4, and 6. Genes related to anti-viral responses (IFIT2, IFIT3, IRF3, MAVS, STING1, TNF) and chemokine trafficking (CCL5, CCL19, CCL21, CCR5, CXCR3) showed variability but were generally expressed at low levels (**Figure 8C**). Particularly noteworthy was cluster 4, which was enriched in SIV-brain. Cluster 4 displayed a gene signature consistent with activated inflammatory macrophages, featuring high expression of CD68, MHC genes, and IL-1β. Upon analyzing the top 10 differentially expressed genes in cluster 4, we identified genes associated with neuroinflammation and neurodegeneration. The adhesion receptor CD44 stood out, having been observed to be upregulated post-brain injury and in microglia/macrophages during neuroinflammation (103). Other genes such as RALGAPA2 and RGCC linked to cognitive impairment and AD were also induced (104, 105) (**Figure 8D**).

**Figure 8.**
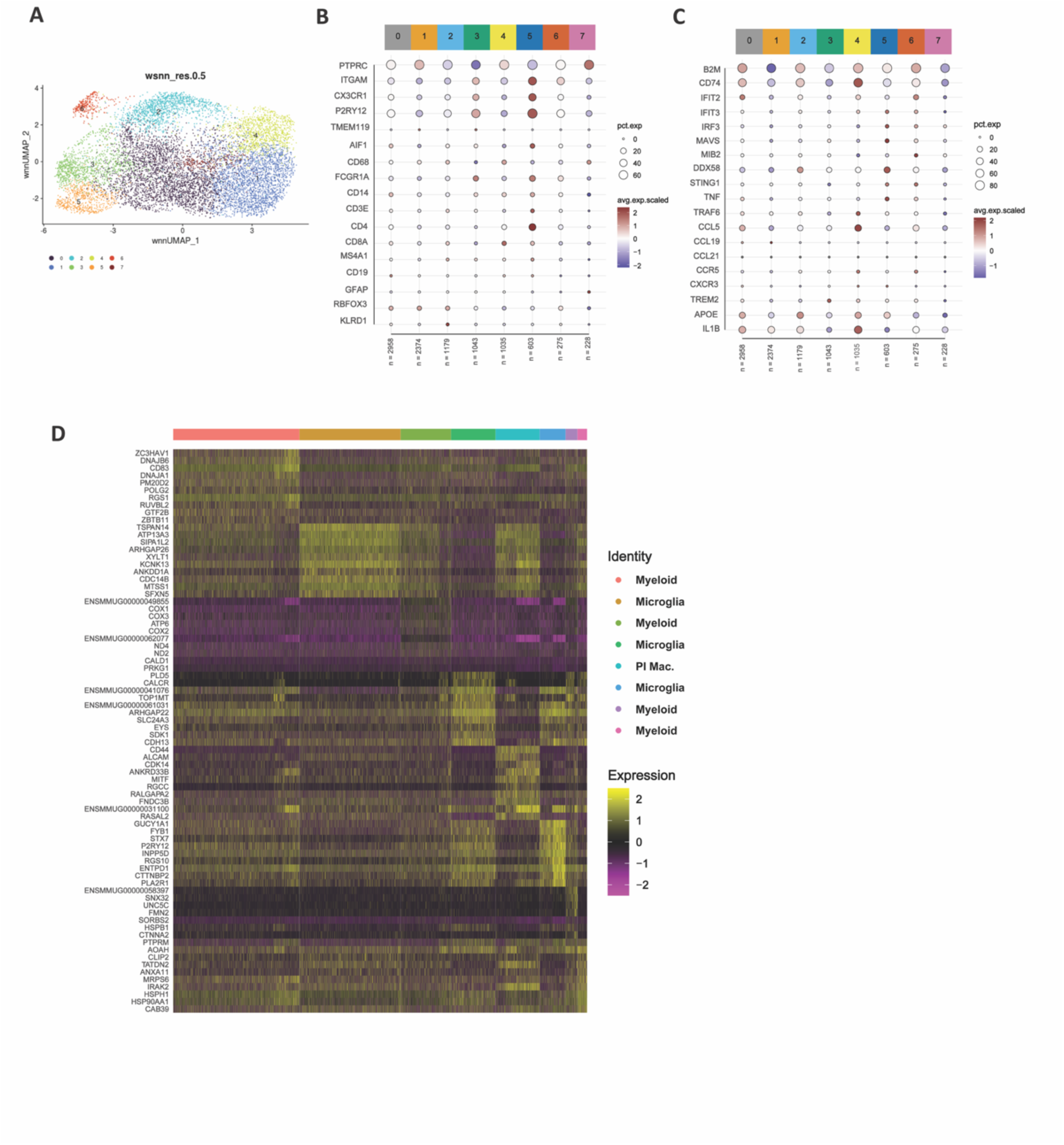
Single cell analysis identifies activated inflammatory macrophage population in SIV infected brain. **(A)** UMAP plot shows cell annotation for myeloid specific gene clusters **(B)** Average expression of common microglia, monocyte, macrophage, and lymphocyte genes across 8 distinct myeloid clusters. **(C)** Average expression of antiviral and inflammatory response genes across 8 distinct myeloid clusters. **(D)** Heatmap of top 10 differentially expressed genes within each cluster.

### CCR7+ CD4 T cell frequencies decreased during SIV-induced neuroinflammation

After uncovering a complex interplay of genes involved in both the initiation and progression of neuroinflammation in response to viral presence in the brain using two complementary methods, we proceeded to examine cellular and soluble markers in the CNS. We examined myeloid populations (microglia [CD11b+CD45lo/int], macrophages, monocytes [CD45+CD14/CD16+]) and lymphoid populations (CD4 and CD8 T cells) from single cell suspensions. Notably, we observed a significant increase in brain monocytes, indicative of active recruitment to CNS during chronic SIV (**Figure 9A**). Correspondingly, there was a significant increase in plasma levels of CCL2, a monocyte chemoattractant, during this infection phase (data not shown). We also investigated microglial activation in the brain and found that overall HLA-DR expression on microglia was not significantly different between controls and chronically infected animals(**Figure 9B**).To explore the potential for active recruitment of monocytes and/or lymphocytes to the brain, we examined a panel of inflammatory analytes. IP-10, a chemoattractant for monocytes and T cells, exhibited a significant increase in the CSF during the chronic infection phase at week 93 post-infection, suggesting ongoing recruitment (**Figure 9C**). While the total CD4 T cell population remained unchanged, the CCR7+ CD4 subset was notably decreased in the brain during chronic SIV infection (**Figure 9D**), an effect not observed in adjacent CNS compartments (**Figure S13**). To determine if the decrease in the CCR7+ CD4 subset was a consequence of virus mediated depletion, we examined expression of chemokine receptors CXCR3 and CCR5 expressed by HIV/SIV target cells. The data showed that, on average, CD4 CD69+ cells in the CNS expressed higher relative levels of CXCR3+CCR5+ arguing against virus mediated depletion of CCR7+ CD4 T cells (**Figure 9E**). In sum, the data show perturbations in neuroimmune homeostasis during SIV infection resulting in decreased CCR7+ CD4 T cells in the brain parenchyma. These findings emphasize the CNS immune surveillance role of CCR7+ CD4 T cells and their potential to counter neuroinflammatory processes during chronic neuroinflammation.

**Figure 9.**
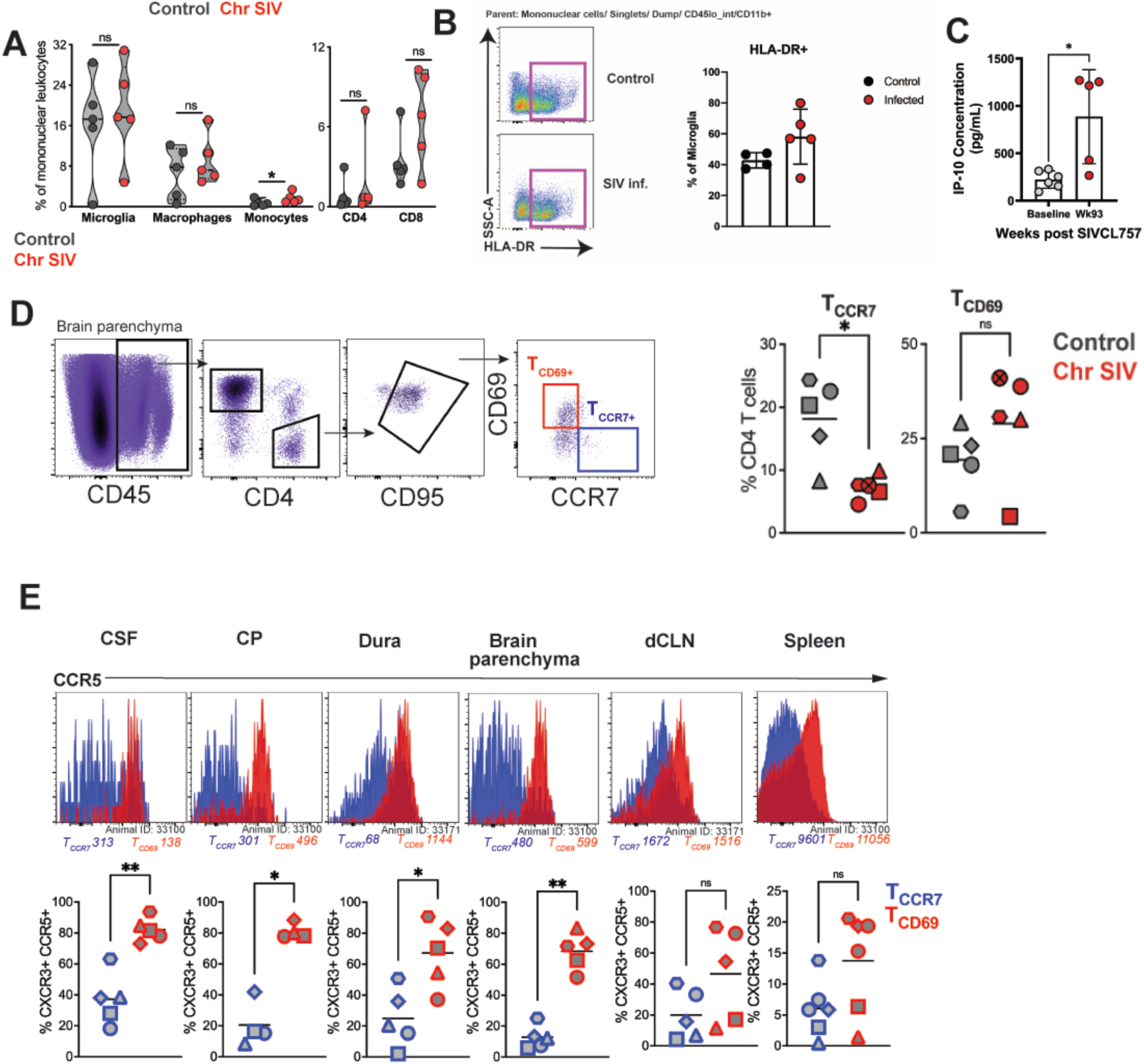
CCR7+ CD4 T cell frequencies decreased during SIV-induced neuroinflammation. **(A)** Frequencies of Myeloid (microglia, macrophage, and monocytes) and Lymphocyte (CD4 and CD8 T cells) cells within the control (black) and chronically infected SIV brain (red). **(B)** Representative flow plot illustrating HLA-DR expression on microglia cells (left) within control (top) and chronically SIV infected (bottom) brain; Frequency of HLA-DR expression (right). **(C)** IP-10 concentration within the rhesus CSF between baseline (grey) and chronic SIV CL757 infection (red; week 93). (**D**) Representative flow plots depict gating strategy for T_CD69+_(red gate) and T_CCR7+_ (blue gate) populations (left); Frequencies of T_CCR7_ and T_CD69_ populations (right) in the control (grey) and chronic SIV CL757 (red) infected brain. (**E**) Histogram plots indicating CCR5 expression and MFI (top) on T_CCR7_ (blue) and T_CD69_ (red); Frequencies of CXCR3+CCR5+ within T_CCR7_ and T_CD69_ across the CSF, Choroid plexus (CP), Dura, Brain Parenchyma, deep cervical lymph node (dcLN) and Spleen.

## DISCUSSION

In this study, we leveraged a multifaceted array of high-dimensional methodologies including single-cell RNA sequencing, integrated single-cell RNA and ATAC sequencing, spatial transcriptomics, and flow cytometry to gain deeper insights into CD4 T cells within the CNS. Our study reveals that T cells present in the non-inflamed CNS parenchyma exhibit diverse differentiation states, characterized by unique chromatin accessibility patterns corresponding to T_CM_ and T_EM_/T_RM_ states. Beyond their presence in the CSF and brain parenchyma, we have also identified a population of T_CM_ cells occupying the skull bone marrow niche, potentially poised to replenish adjacent CNS compartments. Notably, impeding T cell egress from lymph nodes using FTY720 led to a reduction in CCR7+ CD4 T cells within the CSF, suggesting potential migration of T_CM_ from lymph nodes to the CSF, likely through vascular routes. Finally, in a model of chronic HIV infection, we have identified a specific decrease in CCR7+ CD4 T cells in the brain parenchyma.

While the presence of CCR7+ CD4 T cells in the brain challenges established paradigms of memory T cell distribution in non-lymphoid tissues, there is precedence for our observations. For instance, 90% of T cells express CCR7 in human CSF, (12, 106) and CCR7+ T cells populate a variety of other non-lymphoid tissues including the skin, gut, colon, cervix and the rheumatoid synovium (107–109). Moreover, the existence of CCR7+ CD4 T cells in the CNS is not entirely unexpected, given that ligands for CCR7, namely CCL19 and CCL21, are present in human brain lysates and CSF (110–112). In rodent models of experimental autoimmune encephalomyelitis (EAE), endothelial cells constitutively produce CCL19, and inflamed CNS venules express CCL21 (113). In rodent viral neuroinflammation models, there is constitutive CNS expression of CCL19 and CCL21 mRNA, with viral infection enhancing their production and facilitating recruitment of antigen-specific CCR7+ CD8 T cells. Additionally, potential sources of CCL19 include monocytes(114), dendritic cells (115), and microglia(116). Thus, the observed reduction in CCR7+ CD4 T cells with chronic SIV infection may be explained by one of three possibilities: (a) interaction of CCR7 with CCL19; this interaction leading to CCR7 internalization and enhanced adhesion of encephalitogenic T cells to inflamed brain venules is demonstrated in EAE (117, 118), (b) upon encountering cognate antigen on perivascular antigen-presenting cells, CCR7+ CD4 T cells may lose CCR7 expression while gaining tissue homing receptors and effector functions (106, 119, 120) (c) CCR7+ CD4 T cells could be migrating to the draining lymph nodes (such as deep cervical or lumbar lymph nodes) via nasal lymphatics in search of antigens (121). The relative contribution of these processes during chronic viral infections requires further investigation.

Recognizing the potential importance of CCR7+ CD4 T cells in CNS immune surveillance, we hypothesized their presence in the skull bone marrow, given that medullary cavities of bones are established sites for homeostatic proliferation of T_CM_ (122–124). Remarkably, our study unveiled existence of a distinct T_CM_ subset in the skull bone marrow of primates for the first time. The availability of the CCR7 ligand, CCL19, in the CSF (125) raises the possibility that signals conveyed by the CSF may recruit CCR7+ cells from the skull bone marrow to the leptomeningeal space and the brain parenchyma. Recent studies in mice have uncovered a two-way communication between myeloid niches of the skull bone marrow and the meninges (59, 60, 126). For instance, in a mouse stroke model, the skull bone marrow releases a higher number of neutrophils and monocytes to the brain compared to remote long bones (59). Studies in a mouse model of bacterial meningitis demonstrate that CSF interaction with the skull bone marrow niche influences hematopoiesis (127). These findings raise the possibility that mobilization of the lymphoid niche from the skull bone marrow could potentially serve as a local reservoir for replenishing CCR7+ CD4 T cells in perivascular spaces of the brain. Nevertheless, several questions remain unanswered, necessitating further exploration. These include the turnover rate of T_CM_ during homeostasis and infection, sources of IL-7/IL-15 to sustain these populations, the potential contribution of the skull bone marrow compartment during chronic neuroinflammation, and the antigen specificity of the CCR7+ CD4 populations within the CNS.

In addition to CCR7+ CD4 T cells, we show that a distinct subset of CCR7- CD69+ CD4 T cells populate the brain. CD4 T_RM_ have been identified in post-mortem human brain. Characteristically, they express CD69, integrin and chemokine receptors such as CD49a, CXCR6, CCR5, CXCR3, and CX3CR1, along with immune checkpoint molecules like PD-1 and CTLA-4 (48). Notably, CD4 T_RM_ cells diverge from CD8 T_RM_ in that they display lower levels of CD103, with fewer than 10% of brain CD4 T cells exhibiting this marker. The generation of brain T_RM_ following peripheral immunizations raises intriguing questions about the antigen specificities of brain T_RM_ population (128). It also remains to be elucidated whether brain T_RM_ exert a protective role or further amplify neuroinflammation and neurodegeneration during an immune response. A significant aspect to consider is whether distinct T cell subsets, differing in their migration capacities with respect to egress to lymph nodes versus establishing residency, can potentially leverage crucial functions of the immune system to enhance CNS function and sustain immune defense. The sensitivity of neurons to cytotoxic products produced by T cells (129), underscores the adaptive advantage conferred by the presence of cells that can migrate out of the CNS upon activation. Notably, within the SAS, activated CCR7- T cells can enter the brain parenchyma, while quiescent CCR7+ T cells migrate towards lymph nodes in search of antigens (130). The observation of elevated levels of activated T cells within the CSF of people living with HIV who are CSF vRNA+ (131), suggests that such cells could potentially gain access to the CNS parenchyma. These intriguing findings add to the complex landscape of T cell dynamics in the CNS and highlight the need for further exploration to understand their functional implications in both health and disease.

In summary, we demonstrate for the first time that CD4 T cells with T_CM_ features reside within the primate CNS. Taken together these data support a model that the CNS is surveyed by a local CCR7+ T_CM-_like population that provides protection in coordination with CD69+ T_RM_ cells. During chronic viral infection, the T_CM_-like population is perturbed likely due to egress to the draining deep cervical lymph node or differentiation of T_CM_ to T_EM_ in response to local antigen. Further studies defining their migration potential will advance our understanding of neuroimmune surveillance during homeostasis and dysregulation in disease.

### Limitations of the Study

While our study advances our understanding of CD4 T cells in the CNS, it is important to acknowledge its limitations. Although our work provides a deep characterization of T cell differentiation states in the brain, comparative single-cell analyses of T cells across distinct CNS compartments: brain, CSF, skull bone marrow, and deep cervical lymph nodes would be valuable in elucidating the ontogeny and maintenance of CCR7+ CD4 T cells in the brain parenchyma. Additionally, it remains to be determined whether access of CCR7+ CD4 T cells from the SAS or perivascular space to parenchymal regions within the brain is regulated by CCR7-CCL19/CCL21 interactions. Finally, the mechanisms underlying the reduction of CCR7+ CD4 T cells in the brain during chronic inflammation need to be elucidated. Furthermore, exploring whether interventions, such as IL-7/IL-15 treatment, to augment CCR7+ cells in the CNS parenchyma will attenuate chronic neuroinflammation would be of significant value.

## Data and materials availability

RNA-seq dataset is accessible at **GSE221815**.

## MATERIALS AND METHODS

### Rhesus Macaques

Colony bred Rhesus Macaques (*Macaca Mulatta*) were sourced and housed at the California National Primate Research Center (CNPRC) and maintained in accordance with American Association for Accreditation for Laboratory Animal Care (AAALAC) and Animal Welfare Act guidelines. All experimental procedures were approved by the Institutional Animal Care and Use Committee (IACUC) at UC Davis. Animals (total n=47) consisted of both males (n=17) and females (n=30) with ages ranging from 8 months – 29 years. Select animals were utilized for FTY720 treatment studies (n=12) and SIV infection studies (n=5). When available, additional tissues were obtained from uninfected opportunistic medical culls for unrelated conditions (n=9; 4 males, 5 females; ages: 0.8-19.7 [years.months]) to bolster analyses with low sample sizes. Animal details are described in **Table S1 and S2**.

**Table S3** provides details on materials and resources utilized.

### FTY720 Treatment

Fingolimod (FTY720) was obtained from Millipore Sigma and given orally either through mixing with animal feed or by orogastric tube delivery adminsitered during animal sedations during routine sample collections. Animals (n=12, 10 males, 2 females; ages 3.5-5.6 [years.months]) received 30 mg/kg of FTY720 daily over the course of four weeks. Routine collections of both whole blood and CSF were performed on weeks 0 (baseline), 2, 4, 5, and 8 following treatment start and used for subsequent flow cytometric analyses.

### SIV Infection

Animals were infected with a neuropathogenic SIVsm804e-CL757 strain (SIVCL757) donated by Hirsch and colleagues (61), provided at a stock concentration of 1.68 x 10^4^ TCID_50_/mL and stored in liquid nitrogen upon receiving. Prior to infection, viral stocks were thawed at room temperature and diluted 16.7- fold in HBSS to a final volume of 0.5 mL and stored on ice prior to administration. Animals (n=6; 6 females; ages 17.6-20.8 [years.months]) were intravenously infected with 500 TCID_50_ and euthanized at 121 weeks post-infection. During chronic infection, animals were intermittently treated with an anti-retroviral therapy regimen (ART) consisting of nucleoside reverse transcriptase inhibitors, Embicitrabine [40 mg/kg] and Tenofovir Disproxil Fumarate [5.1 mg/kg], and integrase inhibitor, Dolutegravir [2.5 mg/kg]), formulated in a 15% Kleptose aqueous solution and administered as an intravenous daily combination treatment. ART treatments were initiated at either 16 weeks post-infection or when CSF viral loads exceeded 100 vRNA copies/mL and paused once CSF vRNA levels were undetectable. Routine collections of whole blood and CSF were performed at monthly intervals during chronic infection (beyond 12 weeks post-infection).

### Specimen Collection and Processing

Animals were anesthetized with 10mg/kg of ketamine hydrochloride administered intramuscularly for routine collections and with pentobarbital at necropsy. Collection of plasma, serum, and PBMCs were performed as previously described (132). Cerebrospinal fluid (CSF) was collected via the foramen magnum and examined for blood contamination by both visual inspection and Hematsix testing strips (Siemens) in accordance with manufacturer instructions. Cell-free CSF supernatants were collected following centrifugation at 400 rcf for 10 minutes at 4°C. Cell-free CSF was stored at -20°C and utilized for subsequent viral RNA quantification. CSF cell pellets were used immediately for downstream flow cytometric analyses. Lymphoid tissues (draining cervical lymph nodes [dCLN], thoracic lymph node, and spleen) and CNS-associated tissues (brain parenchymal tissues, dura mater, choroid plexus [CP], pituitary gland, and skull bone marrow), following cardiac saline perfusion, were obtained at necropsy and immediately processed for isolation of mononuclear cells.

Brain, dura mater, and skull bone marrow tissues were enzymatically digested in complete media (DMEM supplemented with 10% FBS, 1% penicillin and streptomycin, 2mM L-glutamine) with 250 units/mL of collagenase IV (Worthington Biochemical Corporation), TLCK (N-a-tosyl-L-lysine chloromethyl ketone hydrochloride) trypsin inhibitor, and 5 units/mL of DNase I (Roche Diagnostics) (digestion media) for 45 minutes at 37°C on a shaker at 250 rpm. Digested tissues were then homogenized using a pipette controller fitted with a 10 mL serological pipette, then passed first through a metallic strainer for mechanical dissociation into a petri dish. After the samples were mechanically dissociated, a 180μm nylon strainer and 100µm SMART strainer was utilized to separate fat from the cells. Cells were then washed in media and centrifuged at 400 rcf for 10 minutes. Mononuclear cells were then collected using a 21% and 75% Percoll gradient, washed, counted, and stored in heat-inactivated fetal bovine serum (HI-FBS) with 10% DMSO and stored in liquid nitrogen until subsequent processing.

Spleen, dCLN, and thoracic lymph nodes were mechanically dissociated and incubated in digestion media for 30 minutes at 37^0^C on a shaker at 250 rpm and then passed through both a metallic strainer and a 180 mm nylon strainer. Cells were washed with media and subsequently treated with ACK lysis buffer to eliminate red blood cells. Mononuclear cells were then washed, counted, and stored HI-FBS with 10% DMSO for cryopreservation.

Choroid plexus and pituitary gland tissues were mechanically dissociated and passed through both metallic strainers and 180 mm strainers. Mononuclear cells from choroid plexus tissue were then isolated using a 21% and 75% Percoll gradient. Mononuclear cells from both tissues were subsequently treated with ACK lysis buffer, washed, counted, and stored in HI-FBS with 10% DMSO for cryopreservation.

### Flow Cytometry

Whole blood, CSF and fine needle lymph node aspirates (FNA) were freshly stained and acquired on the same day following collection. Mononuclear cells obtained from necropsy tissues were either freshly stained and acquired the same day or stained following cryopreservation, with identical methods used for all comparisons. For cryopreserved cells, samples were thawed in ice baths and diluted in complete media. Cells were then washed and incubated in complete media with 2 units/mL of DNAse I for 15 minutes at 37°C. Cells were then washed with complete media and counted prior to staining. Whole blood samples were treated with BD FACS Lysing Solution (BD Bioscience) for 10 minutes and washed with 1X FACS buffer (phosphate buffered saline with 1.5mM sodium azide, 2% fetal bovine serum, 10mM EDTA) prior to surface staining. Antibodies for surface staining were prepared in Brilliant Stain Buffer Plus (BD Biosciences) and incubated with cells at 4°C for 30 minutes and washed twice with FACS buffer. For intracellular staining, cells were fixed with Foxp3 Fixation/Permeabilization Buffer following surface staining and incubated at 4°C for 20 minutes and washed with 1X Foxp3 Perm/Wash solution (Invitrogen). Cells were then stained with antibodies for intracellular targets prepared in 1X Foxp3 Perm/Wash solution and incubated at 4°C for 1 hour on a shaker. Next, cells were washed once with 1X Foxp3 Perm/Wash, twice with 1X FACS buffer, and subsequently acquired. Sample acquisition and fluorescence measurements were performed on a BD Bioscience FACSymphony utilizing FACSDiva software (Version: 8. 0.1). Sample compensation, population gating, and analysis was performed using FlowJo (Version 10.8.1)

### Intracellular Cytokine Stimulation Assay

Aliquots of 2 million freshly collected PBMCs and CSF cells were stimulated with PMA/Ionomycin (eBioscience Cell Stimulation Cocktail) and incubated for 1 hour at 37°C. Brefeldin A (BD GolgiPlug) and monensin (BD GolgiStop) were added to cell suspensions and incubated at 37°C for an additional 4 hours. Cells were washed with 0.5 mL of chilled FACS buffer and surface stained in the dark at 4°C for 30 minutes. Cells were then washed with FACS buffer and fixed with Cytofix/Cytoperm solution (BD Bioscience) for 10 minutes at 4°C, subsequently washed with 1X Perm/Wash (BD Bioscience) and treated with intracellular antibody stains prepared in 1X Perm/Wash solution in the dark at 4°C for 30 minutes. Cells were then washed once with 1X Perm/Wash, twice with FACS buffer, and subsequently acquired on a BD FACSymphony.

### FlowSOM

Clustering of cells and construction of a minimum spanning tree of relationships between clusters was conducted using FlowSOM, version 2.4.0 (133) with logicle transformation of data prior to analysis (134). Clustering was performed based on CXCR3, CD95, PD-1, CD69, CCR5, and CCR6 expression among cells, with CCR7 expression used to define groups within metaclusters in subsequent analyses. Other markers used in initial gating steps, or which were ubiquitously expressed/not expressed among cells were excluded from clustering analyses. Numbers of metaclusters were selected dynamically by the FlowSOM algorithm. Data from each panel was analyzed separately with a cluster defined as overrepresented in the CCR7+ or CCR7- group if the corresponding adjusted standardized residual from the chi-square test performed on the table of cluster cell counts by CCR7 status was greater than 3. Analyses were conducted using R version 4.2.1.

### Viral RNA Quantification

Quantification of plasma and CSF viral RNA (vRNA) was performed using a previously described quantitative reverse transcriptase polymerase chain reaction (qRT-PCR) assay for the detection of SIV gag (135). The assay was conducted by the AIDS and Cancer Virus Program at Leidos Biomedical Research Inc., Frederick National Laboratory using methods described by Li et al (136).

### Cell Preparation for Sequencing Studies

Cryopreserved mononuclear cells from rhesus brain and splenic tissues were thawed at room temperature, placed in fresh complete media (For splenic cells: RPMI supplemented with 10% HI-FBS, 1% L-glutamine, 1% penicillin-streptomycin; For brain tissue derived cells: DMEM supplemented with 10% HI-FBS, 1% L-glutamine, 1% penicillin-streptomycin) and treated with 2 units/mL of DNase I (Roche Diagnostics) for 15 minutes at 37°C. Cells were washed in complete media and CD45+ cells isolated using CD45 magnetic bead separation for non-human primates (Miltenyi Biotec CD45 Microbeads non-human primate) in accordance with the manufacturer’s protocol. Enriched CD45+ cells were stained for CD45 and a live dead marker for subsequent flow cytometric sorting. Live CD45+ cells were characterized and quantified on a BD FACSymphony cell analyzer and sorted utilizing a FACS Aria and suspended in DMEM for single cell RNA sequencing studies. Both CD45+ and CD45- cells collected during column enrichment were utilized for single nucleus RNA sequencing and ATAC sequencing studies.

### Single Cell RNA sequencing

Sample barcoding, assembly of gel-beads in emulsion (GEM), GEM reverse transcription, cDNA amplification and cleanup, and library construction were performed according to the Chromium Next GEM single cell 3’ v3.1 protocol from 10X Genomics. Sequencing was performed by SeqMatic LLC on a NovaSeq 6000 platform using S4 200 flow cells with paired end reads ran in four replicates with an average of 111,000 reads per cell.

Sample demultiplexing, generation of FASTQ files, sequence alignment, gene counting, and sample aggregation were performed using the Cellranger pipeline version 7.1.0. Samples that passed data quality control steps (removal of samples with low quality reads, low frequency of mapped reads, low number of reads per cell, high mtRNA signature), were used for subsequent analyses. Sequenced reads were aligned to the Mmul_10 genome reference for Rhesus macaque, and raw count matrices were generated which were used as the input to the Seurat integrated analysis pipeline (Seurat V4.3.0). Quality control was done at the gene and cell level accounting for the median number of genes, and mitochondrial gene percentage using quality control plots. After filtering, graph-based cell clustering with a resolution of 0.5 was performed on statistically significant principal components and visualized using Uniform Manifold Approximation and Projection (UMAP). Cluster identity was assigned by a combination of approaches which includes identifying cluster-specific differentially expressed genes, expert knowledge, canonical list of marker genes, and automated annotations using immune reference atlas through SingleR. T-cell specific clusters were further subset for downstream analysis. Differential gene expression (DEG) analysis of the different cell-types across conditions was performed using the functions from Seurat and were selected at a threshold of (adjusted P-value < 0.05, |log2 FC| > 0.25) based on Benjamini-Hochberg correction. Gene-set enrichment analysis, and functional annotation was implemented through clusterProfiler 4.0, and visualized using custom scripts. All downstream data analysis was performed using R v4.2.0.

### Single nuclei RNA-seq and ATAC

Nuclei were isolated from brain derived live CD45+ and CD45- cells using the Chromium Nuclei Isolation Kit (10X Genomics) in accordance to manufacture instructions. Following isolation, nuclei were stored on ice and used immediately for subsequent library preparation. Barcoded 3’ single cell gene expression library and ATAC library were prepared from single nuclei suspensions using the Chromium Single Cell Multiome kit (10X Genomics) following the manufacturer’s instruction. Briefly, single nuclei suspensions were incubated with transposase to fragment the DNA in open regions of chromatin while adapter sequences were simultaneously added to the ends of the DNA fragments. Transposed nuclei were loaded onto a microfluidic chip and ran in a Chromium Controller instrument to partition individual nuclei into single gel bead forming droplets, or gel beads-in-emulsion (GEMs). Each GEM contained oligonucleotides with a unique 16 base pair 10x Barcode sequence, a poly(dT) sequence to capture polyadenylated mRNA for a gene expression library, and a Spacer sequence that enables barcode attachment to transposed DNA fragments for an ATAC library. GEMs were then incubated to attach the unique barcodes to mRNA and transposed DNA fragments and followed by a reverse transcription reaction, converting mRNA into full-length cDNA. Following cDNA generation, GEMs were disrupted, and pooled fractions recovered, purified, and subjected to a pre-amplification PCR step to fill gaps and ensure maximum recovery of barcoded ATAC and cDNA. Subsequently, the pre-amplified product was used as input for both ATAC library construction and cDNA amplification for gene expression library construction. The cDNA and library fragment size distribution were verified on a Bioanalyzer 2100 (Agilent). The libraries were quantified by fluorometry on a Qubit instrument (LifeTechnologies) and by qPCR with a Kapa Library Quant kit (Kapa Biosystems-Roche) prior to sequencing. The libraries were sequenced on a NovaSeq 6000 sequencer (Illumina) with paired-end 150 bp reads with approximately 35,000 reads pairs per nuclei for gene expression and 25,000 read pairs per nuclei for ATAC libraries.

### Single nuclei RNA-seq and ATAC quantification and statistical analysis

The raw single-cell multiome (ATAC + Gene Expression) sequencing data was pre-processed using Cell Ranger ARC pipeline provided by 10X Genomics. This step involves the demultiplex of cells using cell barcodes, the alignment of reads to Mmul10 reference genome, removal of empty droplets, cell debris and low-quality cells. The filtered gene expression data was imported to Seurat (137)(v4.3.0) for further quality control. Cells were required to have the number of genes expressed in between 250 and 5000, the number of unique transcripts expressed in between 500 and 12000, and the number of ATAC peaks in between 1000 and 70000. Cell doublets were removed using DoubletFinder(138). As a result, the number of cells remaining for downstream analysis are 6375, 12592 and 16742, for samples 32967, 33994 and 33980, respectively.

Gene expression data and chromatin accessibility data were normalized and dimensionality reduced individually. Gene expression data were normalized using LogNormalize method in Seurat with a scale factor of 10000. Cell cycle scores were calculated using previously established cell cycle markers included in Seurat and cell cycle was regressed out during the scaling of the data. Chromatin accessible peak data produced by Cell Ranger was processed using Signac(139) (v1.8.0). It was normalized using term frequency-invese document frequency (TF-IDF) in RunTFIDF function, followed by selecting the top features and then dimension reduced sing singular value decomposition on the TF-IDF matrix. After dimension reduction, the two modalities were integrated using the weighted nearest neighbor method in Seurat. The integrated graph was then used for UMAP visualization and clustering. Cell type identification was carried out on clusters generated at resolution 2.25, using the R package scType(140). Differential expression analysis was done using linear model in limma(141), adjusting for cell cycle scores and the number of genes expressed.

### Spatial Transcriptomics Profiling

Paraformaldehyde-fixed, paraffin-embedded hippocampal brain tissue from necropsied non-human primates (uninfected control and SIV infected) was profiled using GeoMx^®^ DSP (142). 5 µm tissue sections were prepared according to manufacturer’s recommendations for GeoMx-NGS RNA BOND RX slide preparation (manual no. MAN-10131-02). Deparaffinization, rehydration, heat-induced epitope retrieval (for 15 minutes at 100°C), and enzymatic digestion (0.1 μg/mL proteinase K for 15 minutes at 37°C) were carried on the Leica BOND-RX. Tissues were incubated with 10% neutral buffered formalin for 5 minutes and for 10 minutes with NBF Stop buffer. Tissue morphology was visualized using fluorescent antibodies specific to lymphocyte and neuron specific molecular markers (anti-CD45 [Novus], anti-CD3 [Primary, Bio-Rad], Secondary anti-Rat IgG, [ThermoFisher], and anti-NeuN [Millipore Sigma], and nucleic acid stain Cyto83 [ThermoFisher]). Twelve regions of interest (ROIs) of 660 x 785 µm geometric shapes (squares) were created and localized to brain blood vessels and neuron rich areas. After ROIs were selected, UV light was utilized to release and collect oligonucleotides from each ROI. During PCR, Illumina i5 and i7 dual-indexing primers were added to each area of illumination (AOI). Library concentration was measured using a Qubit fluorometer (Thermo Fisher Scientific), and quality was assessed using a Bioanalyzer (Agilent). The Illumina NextSeq 2000 was used for sequencing, and the resulting FASTQ files were then mapped to the Hs_R_NGS_WTA_v1.0 reference (Nanostring) using the NanoString GeoMx NGS pipeline v2.1 to generate raw count data for each target probe in every AOI. The raw counts were processed using the same NanoString GeoMx NGS pipeline and converted to digital count conversion (DCC) files.

The raw count data (DCC files) were further processed using Geomxtools(143) R package (Bioconductor version 3.2.0, Nanostring, Seattle, WA, USA). Data were quality controlled per individual AOI. AOIs were excluded from the dataset if they met any of the following conditions: less than 80% reads aligned to the reference, less than 40% sequencing saturation, or less than 1000 raw reads. Limit of quantification (LoQ) is calculated for raw data based on the distribution of the negative control probes (“NegProbe”) and is used as an estimate for the quantifiable limit of gene expression per AOI (144). A gene is considered detected if its expression is above the LoQ for that AOI. Genes were included in the analysis if they were detected above the LoQ in > 5% of AOIs. Then, the data was normalized using the third quantile (Q3) to account for differences in cellularity and ROI size. Following filtering and normalization we were able to analyze 9,605 genes within 20 segments spread across two animals (Control and SIV-infected). The Linear Mixed Model (LMM) was used to calculate the significant differences between the two groups, and genes were considered significantly expressed when adj.p < 0.1 (677 DEGs). The differentially expressed genes were used to create heatmaps of selected KEGG pathways. The heatmaps were created using ComplexHeatmap (version 2.13.1) R package [citation: Gu, Z. (2022) Complex Heatmap Visualization. iMeta.].

### Immunohistochemistry

De-identified formalin-fixed paraffin embedded human hippocampal tissues from non-demented patients (n=3) and a patient with gliobastoma (n=1) were obtained from the Netherlands Brain Bank and human tonsil tissues (n=1) was obtained from the UC Davis Cancer Center Repository. Rabbit polyclonal anti-CD3 (Agilent, dilution at 1:75), mouse monoclonal anti-CD4 (Novus, dilution at 1:150), rabbit polyclonal anti-CD45 (dilution at 1:20), rabbit monoclonal anti-IBA1 (Invitrogen, dilution 1:100), rabbit polyclonal anti-CD11b (Invitrogen, dilution 1:100), and rabbit monoclonal anti-NeuN (Abcam) were used for subsequent immunohistochemical staining. All 4 µm paraffin sections were subjected to a heat antigen retrieval step before application of primary antibodies by treating slides with AR-10 (Biogenex) for 2 minutes at 125°C in a Digital Decloaking Chamber (Biocare), followed by cooling to 90°C for 10 minutes, or by treating slides with H-3300 (Vector) for 20 minutes at 100°C. Following primary staining, samples were incubated with anti-mouse and anti-rabbit EnVision+ system secondary antibodies (Agilent), followed by treatment with AEC chromogen (Agilent) and counterstained with Gill’s hematoxylin I (StatLab). Primary antibodies were replaced by mouse or rabbit isotype controls and ran with each staining series as negative controls. Slides were visualized with a Zeiss Imager Z1 (Carl Zeiss) and images captured using a Zeiss Axiocam (Carl Zeiss).

### Cerebrospinal Fluid and Serum Biochemistries

Animal CSF and serum chemistries were quantified using a Piccolo Xpress Chemistry Analyzer (Abbott) with Piccolo BioChemistry Plus disks in accordance with manufacturer’s instructions. Chemistry panel analytes include alanine aminotransferase (ALT), albumin, alkaline phosphatase (ALP), aspartate aminotransferase (AST), gamma glutamyltransferase (GGT), lactate dehydrogenase (LDH), C-reactive protein (CRP), creatine kinase, creatine, glucose, total protein, calcium, blood urea nitrogen (BUN), cholesterol, triglycerides, bilirubin, sodium, potassium, chloride, carbon dioxide, and phosphorous.

### Statistical Analyses

Wilcoxon signed rank test were used for paired analyses (i.e., longitudinal and within group comparisons). Mann-Whitney U-test were used for unpaired comparison between animal cohorts/treatment groups. Test were performed in GraphPad Prism (Version 9.5.1) with significance values denoted as follows: * p < 0.05, ** p < 0.01, *** p < 0.001, **** p < 0.0001.

## Supporting Information

**Table S1. NHP cohort for chronic SIVCL757 study**

**Table S2. NHP cohort used for tissue assessment during necropsies/medculls Table S3. Key reagents and resources**

**Figure S1. Distribution of CD4 and CD8 T cells in brain parenchyma and its border tissues**, **Related to** Figure 1 (**A**) Illustrates CD4 and CD8 T cell frequencies (gated on CD3+ cells) in various tissues as indicated. CSF, cerebrospinal fluid; CP, choroid plexus; Dura, Dura mater; Pit, Pituitary; dCLN, deep cervical lymph nodes, Th LN, thoracic lymph nodes. (**B**) bar graphs show % CD3+ T lymphocytes and CD4: CD8 ratios across CNS and lymphoid tissues.

**Figure S2. Single-cell transcriptomic analyses of CD45+ immune cells reveal presence of core T cell molecular programs in brain, related to** Figure 1. (**A**) lists cell numbers sequenced for each sample. (**B-D**) Mean, median reads per cell and UMI counts per cell. (**E**) quality metrics on each sample. (**F**) Expression profile of PTPRC shown in UMAPs and violin plots. (**G**) Expression profile of CD3δ subunit shown in UMAPs and violin plots. (**H**) UMAP plots show cell type annotation for B cells and T cells across tissue types.

**Figure S3. Single-cell transcriptomic analyses of CD45+ immune cells reveal presence of core T cell molecular programs in brain, related to** Figure 1. (**A**) UMAP of scRNA seq transcriptome profiles color coded according to assigned cell types by tissue- brain (2137 cells) and spleen (1569 cells). (**B**) UMAP of T cell subclusters color coded according to assigned cell types by tissue - brain (1619 cells) and spleen (1159 cells). (**C**) Expression of top marker genes in each T cell cluster. (**D-E**) heat maps of gene expression within clusters demonstrate differential expression across clusters.

**Figure S4. Single cell regulatory landscape of T cells and brain resident microglia, related to** Figure 2. (**A**) lists cell numbers sequenced for each sample. (**B-D**) Mean, median reads per cell and UMI counts per cell. (**E**) Cell counts for each cluster shown per sample. (**F**) Expression profile of PTPRC shown in UMAPs and violin plots. Scatter plots show proportion and intensity of gene expression comparing (**G**) Microglia to T cells. (**H**) Macrophage to T cells. (**I-K**) Genomic regions showing snATAC-seq tracks of chromatin accessibility of STAT4, IFNγ, and BCL-2

**Figure S5. CCR7+ CD4 T cells in the CNS share phenotypic features with T_CM_ in blood and lymph nodes, related to** Figure 3**. (A)** Representative flow plots identifying CD3 T cells, CD20 B cells, CD4 T cells, CD8 T cells and Monocytes (Left) and their frequencies (Right) in the CSF, Lymph node Fine Needle Aspirate (LN FNA), and Blood of healthy rhesus macaques. CSF: n=10; LN-FNA: n =12; Blood: n=12 **(B)** Representative flow plots identifying expression of CD28 and CD95 on CD4 and CD8 T cells (Left) and their frequencies (Right) from the CSF, LN (FNA), and Blood. **(C)** Representative flow plots identifying CCR7 expression on CD28+ CD4 and CD28+ CD8 T cells (Left) and their frequencies in the CSF, Lymph node Fine Needle Aspirate (LN FNA), and Blood (D)

**Figure S6. CCR7^+^ CD4 T cells in the CNS share phenotypic features with T_CM_ in blood and lymph nodes, related to** Figure 3**. (A)** Minimal spanning tree constructed using the FlowSOM (self-organizing map) tool to map phenotypic relationships between live CD4 T cell isolated from healthy rhesus CSF. Star chart wedges denote expression of each indiciated marker in cluster. Nodes indicate cellular cluster which are grouped into 5 larger Metaclusters indicated by color (metacluster 1: pink; metacluster 2; yellow; metacluster 3: green; metacluster 4: blue; metacluster 5: purple) which share similar phenotypes. Connections indicate phenotypically related cellular clusters. Nodes are represented as pie charts representing relative expression of chemokine and activation markers (CCR6, CCR5, CD69, CXCR3, CD95, PD-1) within a given cluster. **(B)** Minimal spanning trees highlighting in blue, clusters that contain CCR7- (top) and CCR7+ (bottom) CD4 T cell populations.

**Figure S7. CCR7^+^ CD4 T cells in the CNS share phenotypic features with T_CM_ in blood and lymph nodes, related to** Figure 3. Representative tSNE plot illustrating CD69, BCL2, CD69 (column 1), CCR5, EOMES, CD127 (Column 2), HLA-DR, ICOS (Column 3), and CXCR3, Foxp3, PD-1 (Column 4) expression on CD4+CD28+CD95+ CCR7-/+ T cells in the CSF, Blood and PBMCs; frequencies for each population are expressed to the right of the tSNE plots.

**Figure S8. FTY720 mediated sequestration of CD4 T_CM_ in lymphoid tissues decreases CCR7^+^ CD4 T cell frequencies in CSF, related to** Figure 4.(**A**) Frequencies of Naïve CD4 T cells and Naive CD8 T cells in the blood over the course of the study (**B**) Representative longitudinal flow plots indicating CD28 and CD95 expression on CD4 T cells from the blood (top row) or the CSF (bottom row) (Left); Frequencies of CD28+ memory CD8 T cells, Central Memory CD8 T cells, CCR7+CD28+ memory CD8 T cells, and Effector CD8 T cells in the blood and CSF over the course of the study

**Figure S9. CCR7+ CD4 T cells in CNS exhibit functional T_CM_ features and reside within skull bone marrow, related to** Figure 5. Frequencies of Lineage (CD20/CD3/Dead) negative CD45+ cells (gray), MHC-II+ (DR) (pink), CD11b+ (blue), and DR-CD11b- cells within the choroid plexus (CP)/dura mater (Dura), brain, skull bone marrow (Sk BM), and draining cervical lymph node (dCLN). Box and whisker plots indicate medians with quartiles.

**Figure S10. CCR7+ CD4 T cells in CNS exhibit functional T_CM_ features and reside within skull bone marrow, related to** Figure 5. Frequencies of Lineage (CD20/CD3/Dead) negative CD45+ cells (gray), MHC-II+ (DR) (pink), CD11b+ (blue), and DR-CD11b- cells within the choroid plexus (CP)/dura mater (Dura), brain, skull bone marrow (Sk BM), and draining cervical lymph node (dCLN). Box and whisker plots indicate medians with quartiles.

**Figure S11. Longitudinal Plasma and CSF viral loads, related to** Figure 6. Kinetics of vRNA in Plasma (red line) and CSF (blue) following SIVCL757 infection. Green bars indicate timing of antiretroviral therapy.

**Figure S12. Spatial Transcriptomics reveals genes consistent with T_CM_ cell states within the perivascular regions of rhesus Hippocampus, related to** Figure 7. T lymphocytes are localized to blood vessel in the non-demented human brain. H&E with immunohistochemical staining of lymphocyte specific markers (CD3, CD4), myeloid/leukocyte specific markers (CD11b, CD45, IBA1) and neuron-specific markers (NeuN) from paraffin embedded human tonsil and brain tissues derived from either a patient with glioblastoma (Sample ID: GBM-01) or non-demented patients (Sample IDs: 90-128, 06-080, 98-0189, 06-080). Tissue/patient samples are organized by columns and immunohistochemical markers by row.

**Figure S13. Preferential depletion of parenchymal CCR7+ CD4 T cells in chronic SIV infection, related to** Figure 9. CCR7+ memory T cells in control (grey) and Chronic SIV infected (red) in the CSF, Dura, DCLN, and Spleen (left to right).

## Acknowledgements

The authors would like to extend their sincere gratitude to several individuals whose contributions have been crucial to the successful execution of these studies. Special appreciation is expressed to Wilhelm Von Morgenland and Miles Christensen, along with the CNPRC SAIDS team, for their coordination of macaque studies, animal care, and ongoing support. The authors also wish to acknowledge the dedicated efforts of the CNPRC Veterinary Staff in ensuring the well-being of the animals and their assistance throughout the study. Brian Schmidt’s exceptional technical assistance in the preparation of ART and acquisition of flow cytometry data is acknowledged. The authors would also like to convey gratitude to Jennifer Watanabe and members of Koen Von Rompay’s laboratory for their valuable technical support during necropsies. The authors extend gratitude to Drs. Vanessa Hirsch and Cheri Lee for providing the SIVCL757 virus. Appreciation is also expressed to Dr. Paul Luciw and Dr. Rompay for their valuable intellectual contributions. Furthermore, the authors extend their gratitude to the Netherlands Brain Bank for providing the valuable tissue specimens that were essential for this research. The authors acknowledge funding support from NIAID, NIH K01OD023034 (SSI); NIA, RF1AG06001 (SSI, JHM); NIAID, NIH R56AI150409 (SSI) and support with federal funds from the National Cancer Institute, National Institutes of Health under contract number HHSN261201500003I and NCI contract 75N91019D00024 (JDL).

