## Supplemental Figures for "CCR7+ CD4 T Cell Immunosurveillance Disrupted in Chronic SIV-Induced Neuroinflammation in Rhesus Brain"

### Figure S1.

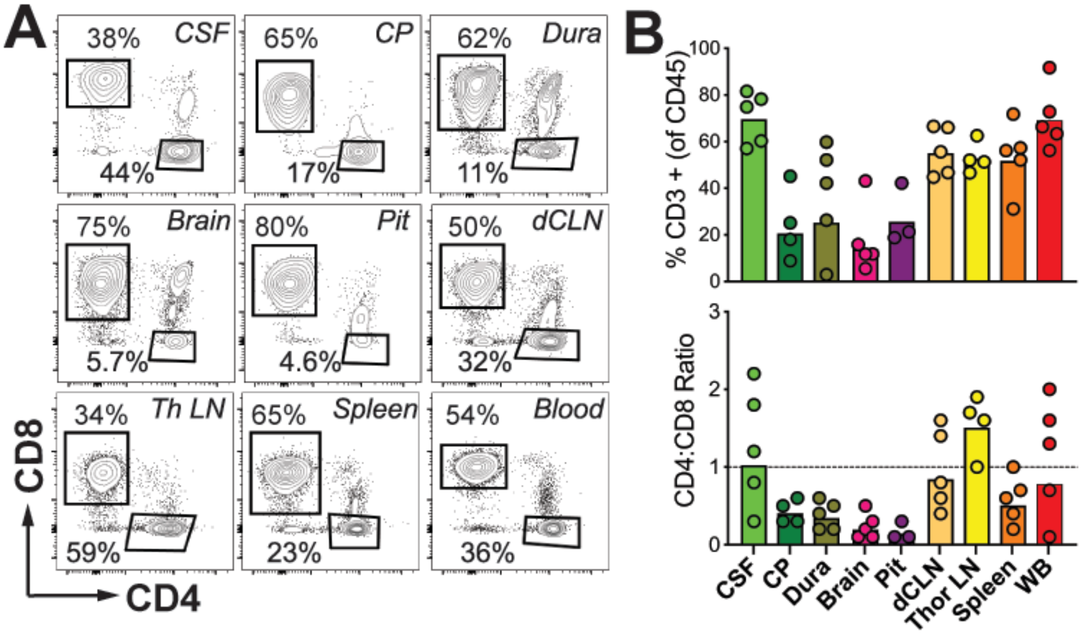


### Figure S2.

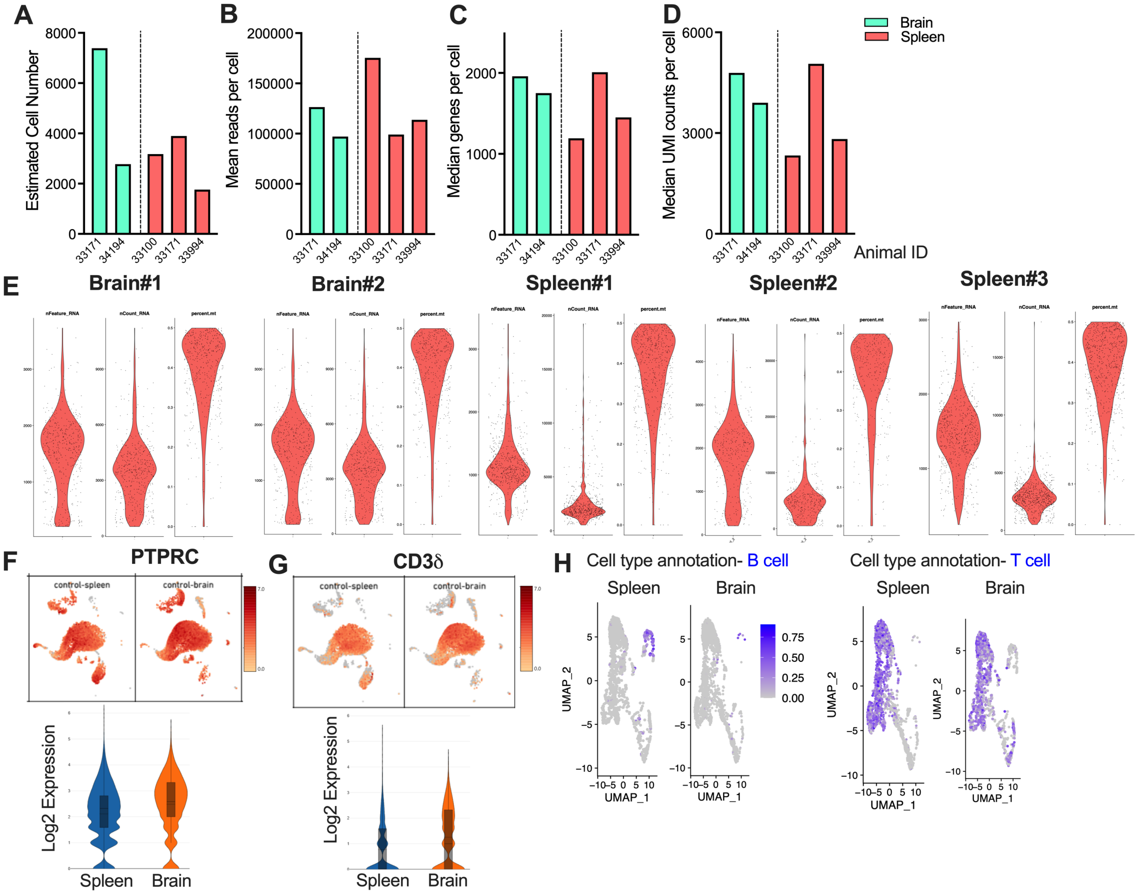

**Figure S3.**

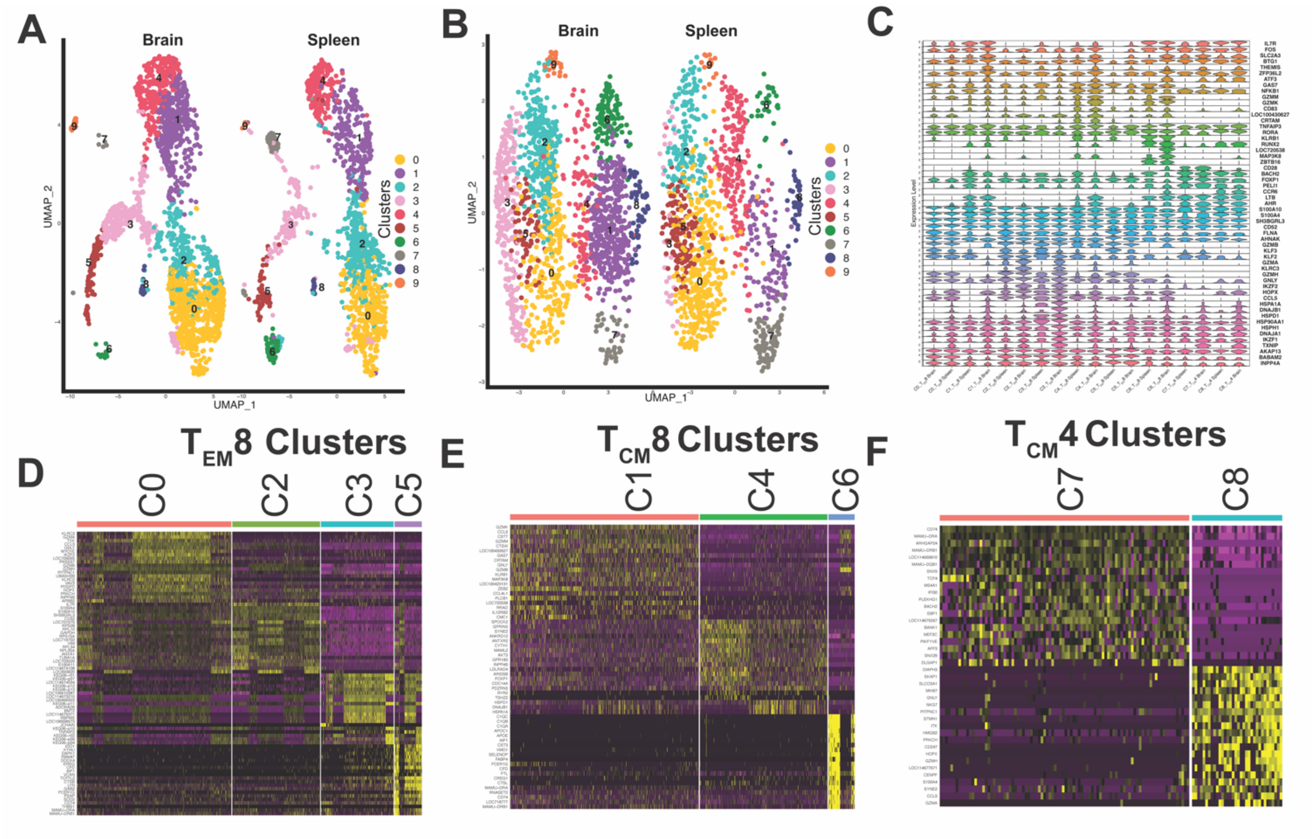

A
Brain
Spleen
UMAP_2
UMAP_1
Clusters
B
C
Expression Level
D
TEM8 Clusters
C0
C2
C3
C5
E
TCM8 Clusters
C1
C4
C6
F
TCM4 Clusters
C7
C8

**Figure S4.**

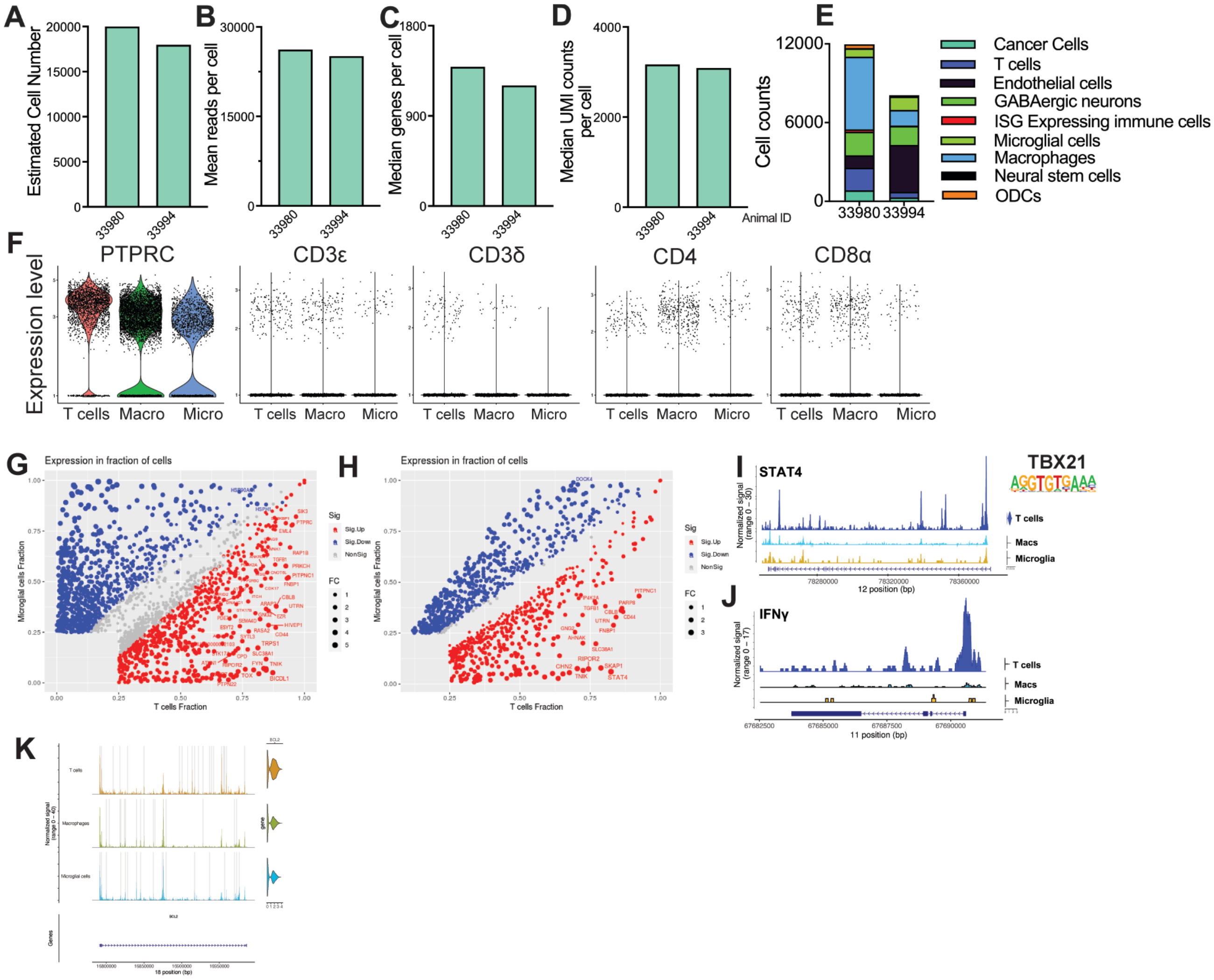

A
Estimated Cell Number
20000
15000
10000
5000
0
33980
33994
B
Mean reads per cell
30000
15000
0
C
Median genes per cell
1800
900
D
Median UMI counts per cell
4000
2000
Animal ID
E
Cell counts
12000
6000
Cancer Cells
T cells
Endothelial cells
GABAergic neurons
ISG Expressing immune cells
Microglial cells
Macrophages
Neural stem cells
ODCs
F
Expression level
PTPRC
CD3ε
CD3δ
CD4
CD8α
T cells
Macro
Micro
G
Expression in fraction of cells
Microglial cells Fraction
T cells Fraction
Sig
Sig.Up
Sig.Down
NonSig
FC
H
I
STAT4
TBX21
T cells
Macs
Microglia
12 position (bp)
J
IFNγ
11 position (bp)
K
Macrophages
Microglial cells
Genes
18 position (bp)

**Figure S5.**

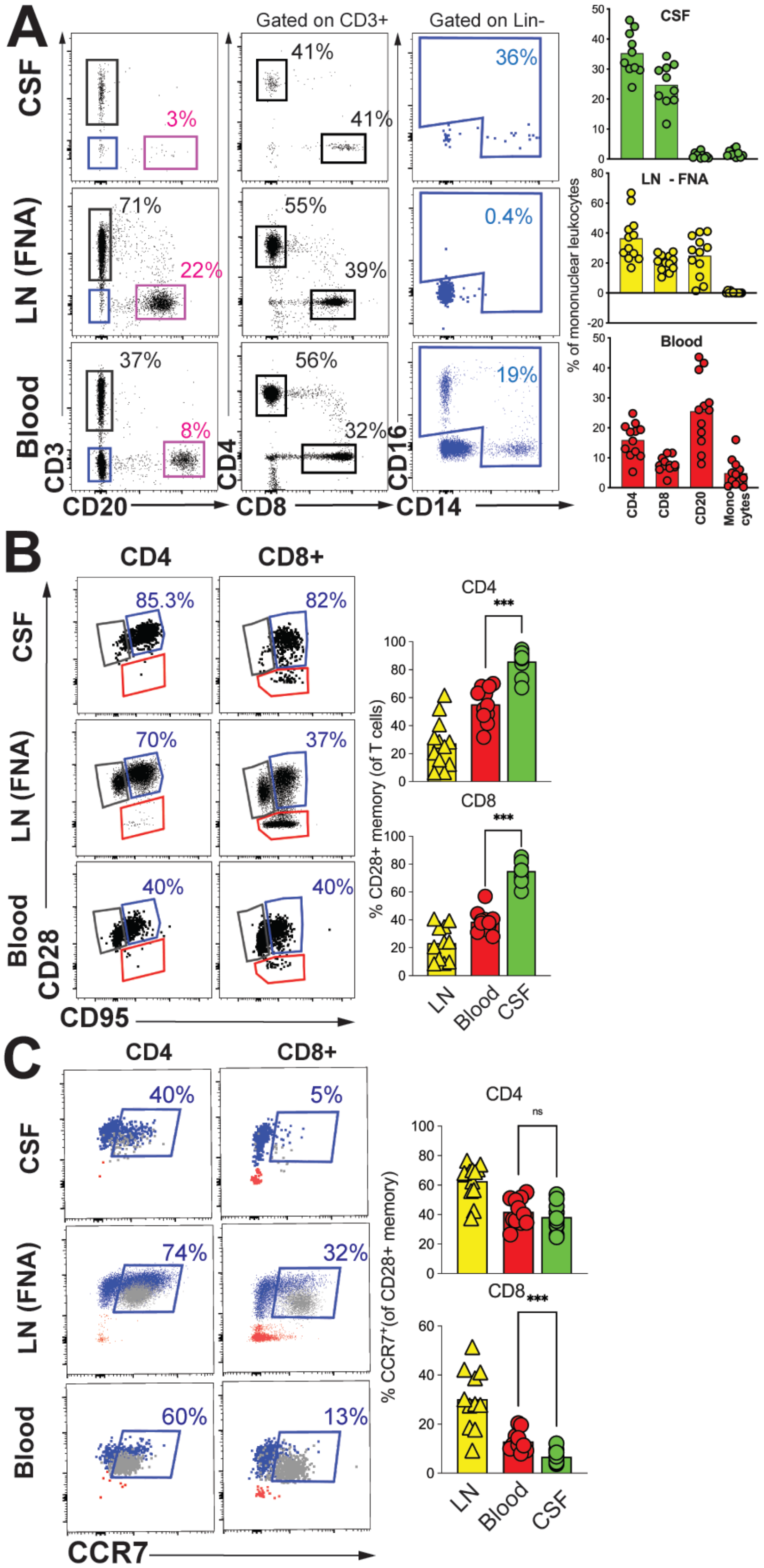

A
CSF
LN (FNA)
Blood
Gated on CD3+
Gated on Lin-
3%
22%
8%
71%
37%
41%
41%
55%
39%
56%
32%
36%
0.4%
19%
CD3
CD20
CD4
CD8
CD16
CD14
CSF
LN - FNA
Blood
% of mononuclear leukocytes
CD4
CD8
CD20
Mono cytes
B
CD4
CD8+
85.3%
82%
70%
37%
40%
40%
CD28
CD95
CD4
CD8
***
% CD28+ memory (of T cells)
LN
Blood
CSF
C
CD4
CD8+
40%
5%
74%
32%
60%
13%
CCR7
CD4
ns
CD8
***
% CCR7+(of CD28+ memory)
LN
Blood
CSF

**Figure S6.**

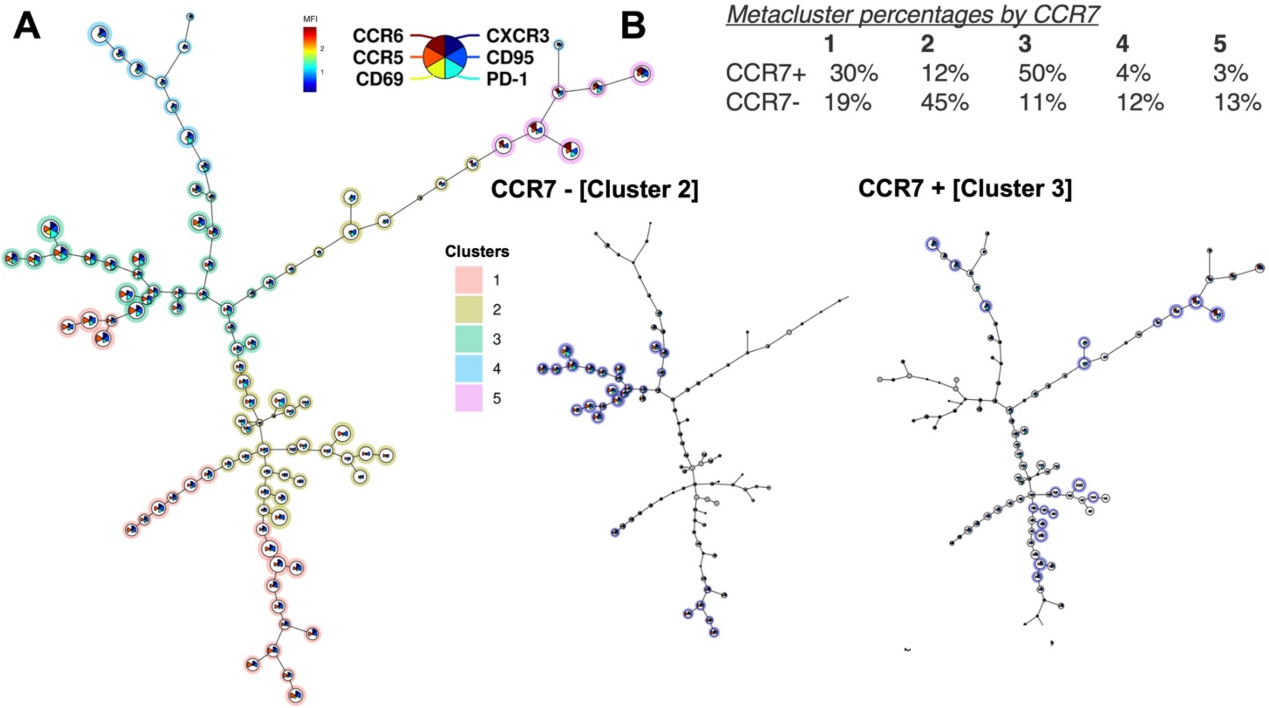


*Metacluster percentages by CCR7*

| | 1 | 2 | 3 | 4 | 5 |
|---|---|---|---|---|---|
| CCR7+ | 30% | 12% | 50% | 4% | 3% |
| CCR7- | 19% | 45% | 11% | 12% | 13% |

Figure S7.

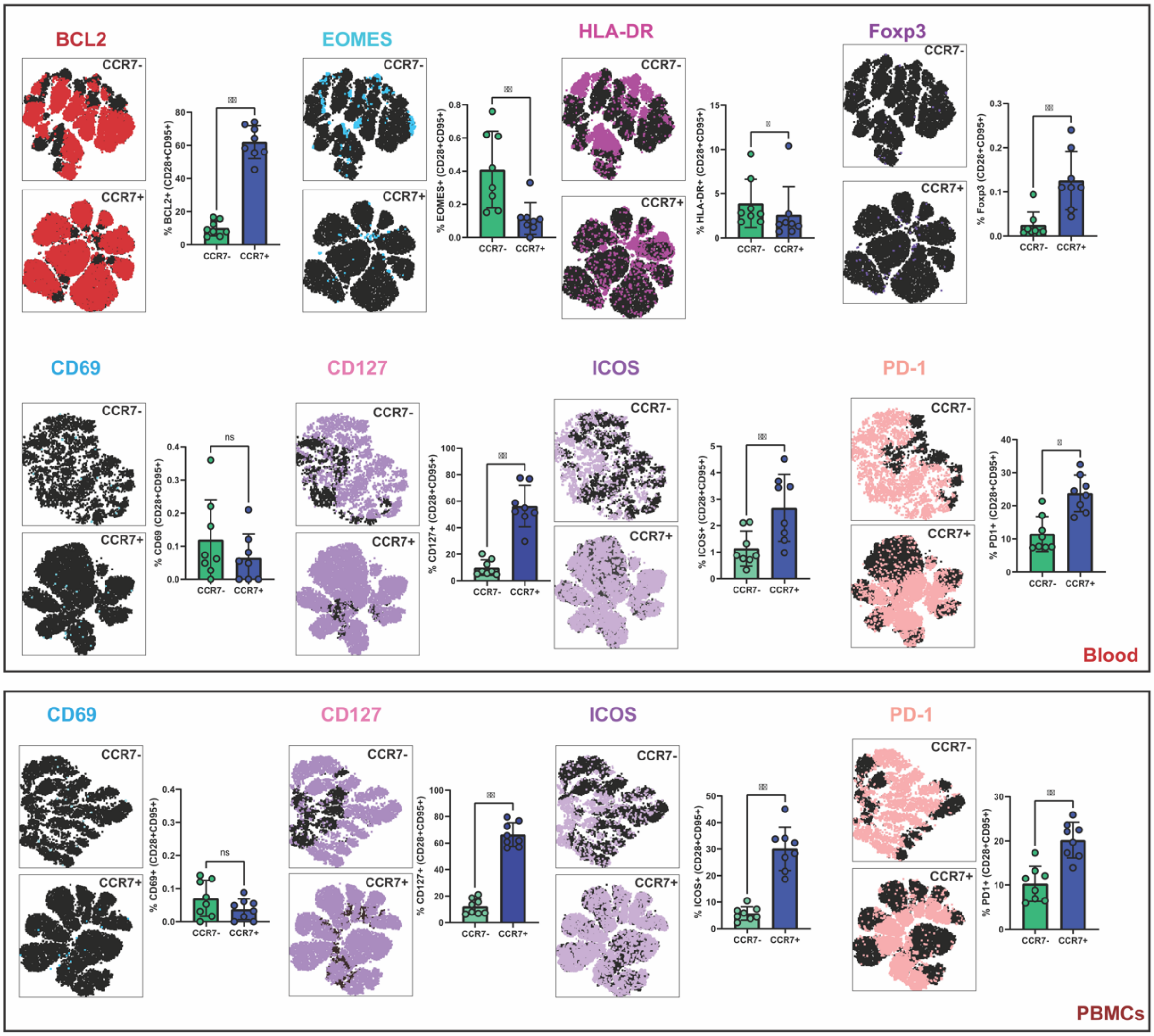


Figure S8.

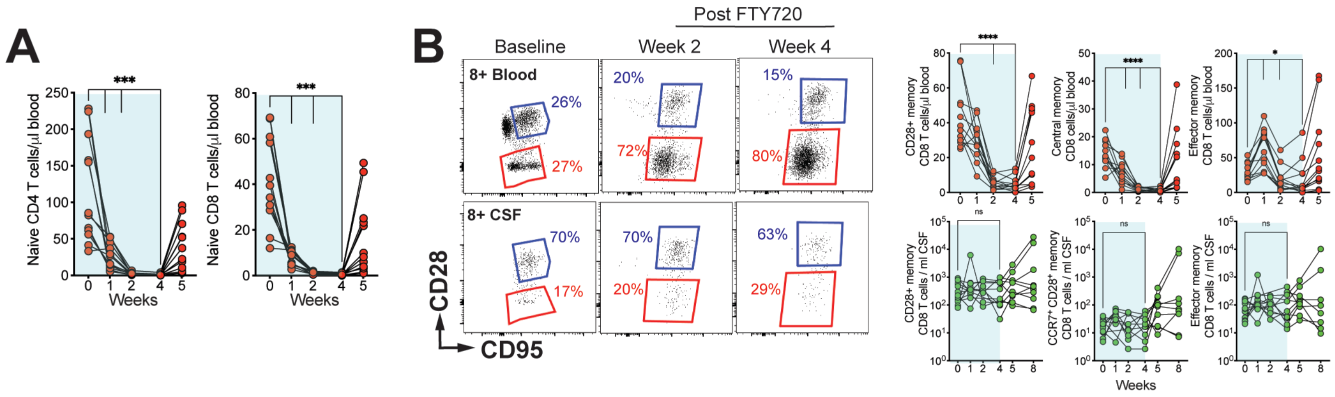

**Figure S9.**

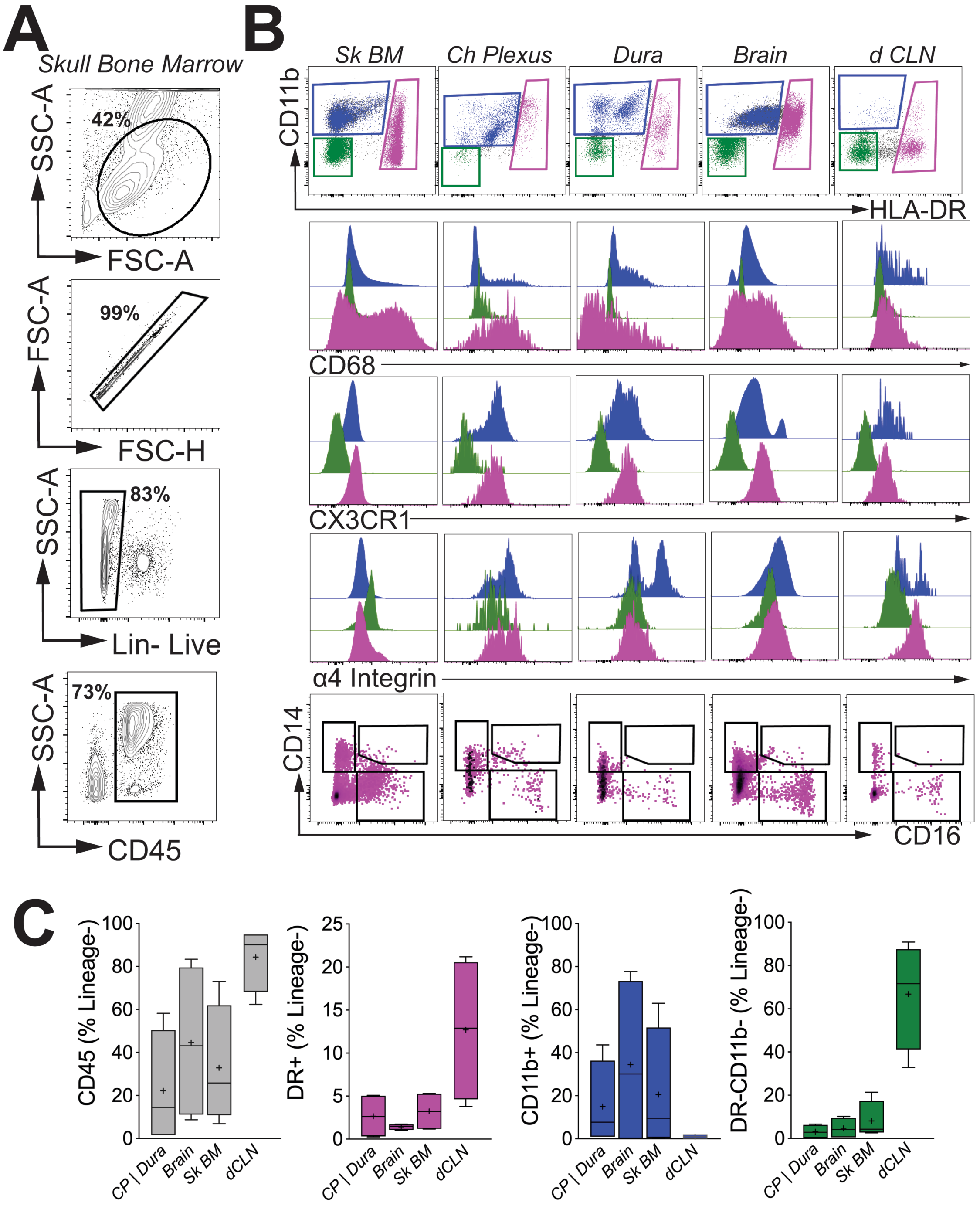
A
Skull Bone Marrow
SSC-A
FSC-A
42%
FSC-H
99%
Lin- Live
83%
CD45
73%
B
Sk BM
Ch Plexus
Dura
Brain
d CLN
CD11b
HLA-DR
CD68
CX3CR1
α4 Integrin
CD14
CD16
C
CD45 (% Lineage-)
DR+ (% Lineage-)
CD11b+ (% Lineage-)
DR-CD11b- (% Lineage-)
CP | Dura
Brain
Sk BM
dCLN

**Figure S10.**

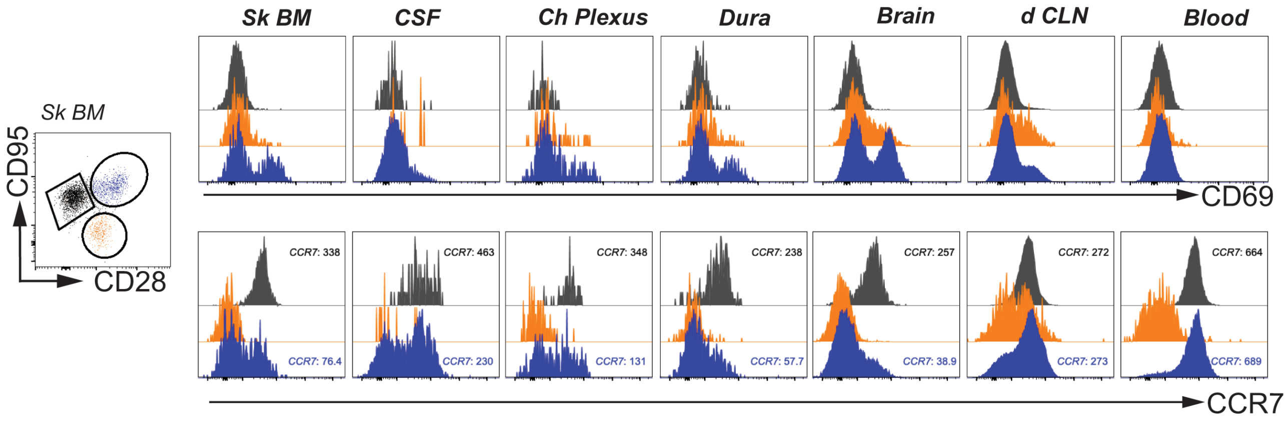


**Figure S11.**

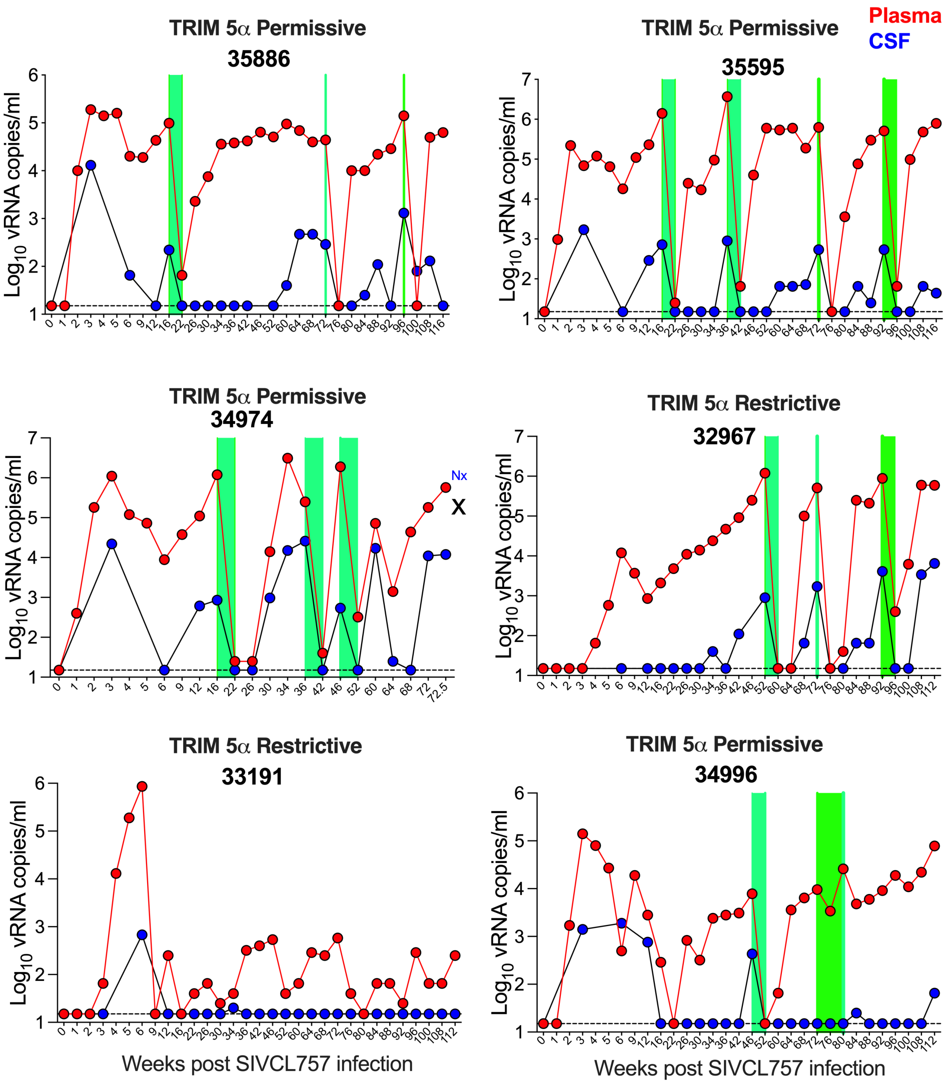

**Figure S12.**

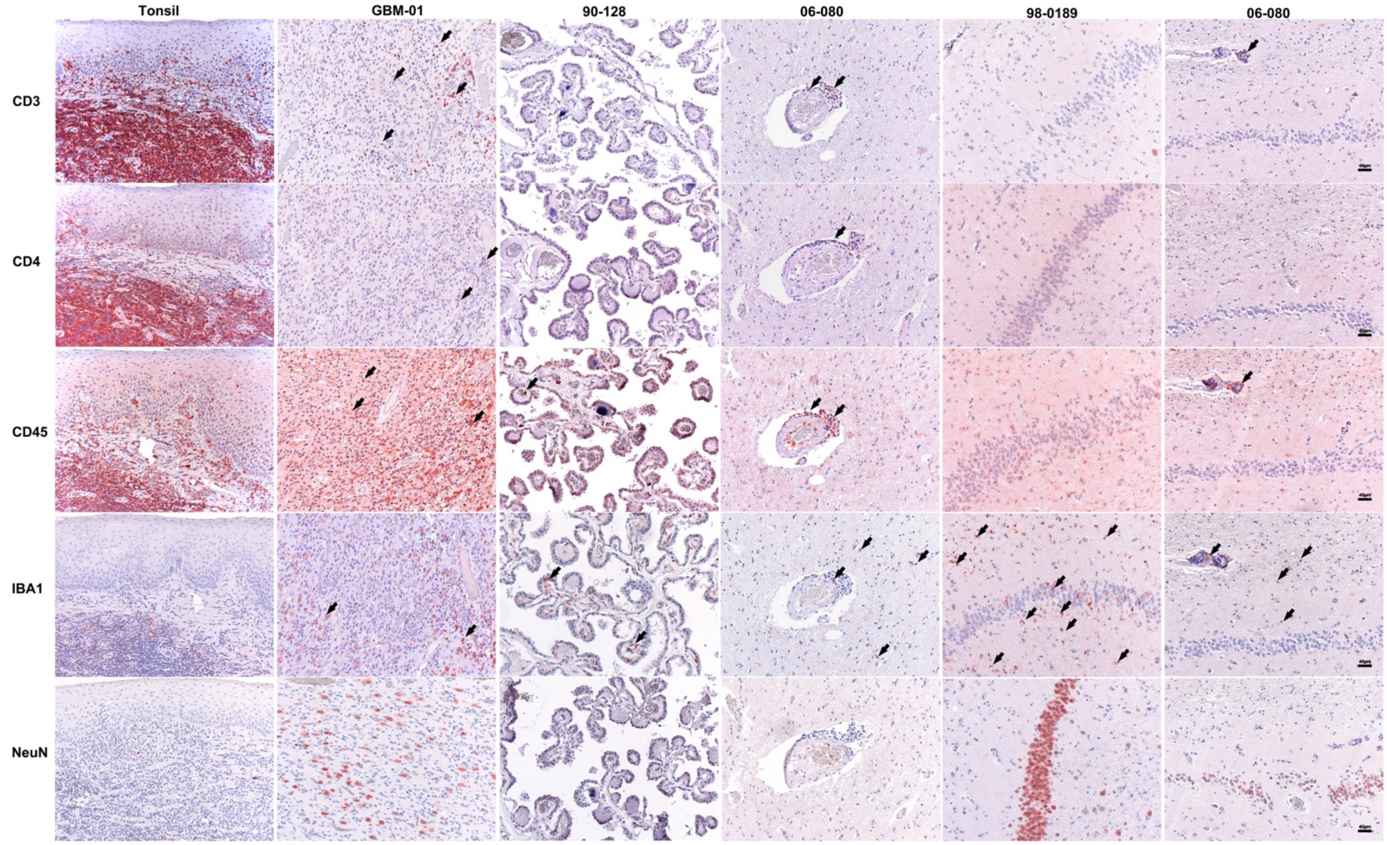


**Figure S13.**

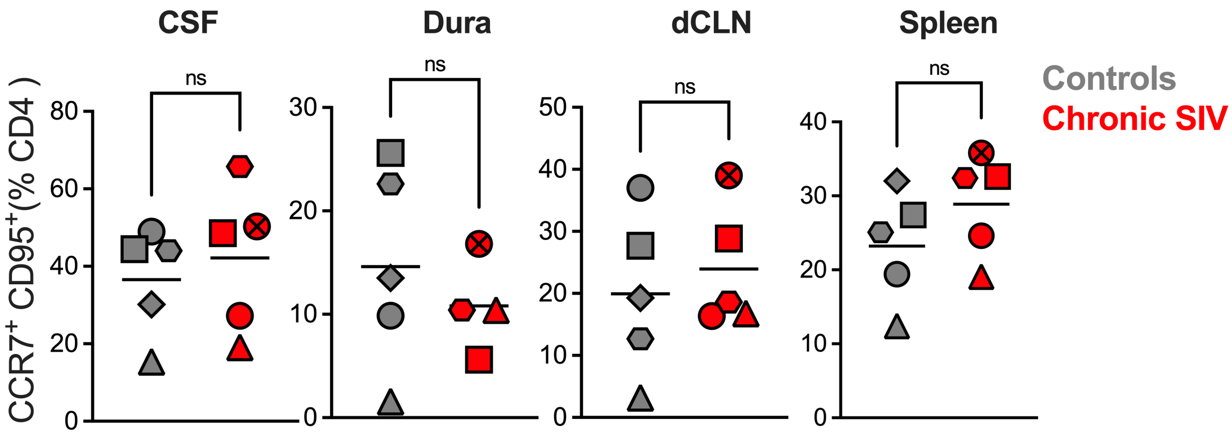
