## Supplemental Tables for "CCR7+ CD4 T Cell Immunosurveillance Disrupted in Chronic SIV-Induced Neuroinflammation in Rhesus Brain"

#### S1 Table. NHP cohort for chronic SIV study (SIVCL757)

| Animal ID | Infection status | Age at Euthanasia(years:months:day) | Sex | Weight (kg) | TRIM5 | ART regimen |
|---|---|---|---|---|---|---|
| 33100 | Uninfected | 20:07:09 | **F** | **12.32** | TFP/Q | N/A |
| 33171 | Uninfected | 20:06:27 | F | 9.27 | TFP/TFP | N/A |
| 33980 | Uninfected | 19:06:26 | F | 9.6 | TFP/TFP | N/A |
| 33994 | Uninfected | 19:06:16 | F | 10.31 | TFP/TFP | N/A |
| 34194 | Uninfected | 19:06:16 | F | 11.84 | TFP/TFP | N/A |
| 35886 | SIVCL757 | 17:05:27 | F | 9.11 | TFP/Q | FTC/TDF/DTG |
| 35595 | SIVCL757 | 17:07:17 | F | 11.9 | Q/CypA | FTC/TDF/DTG |
| 34974 | SIVCL757 | 17:08:22 | F | 8.94 | TFP/Q | FTC/TDF/DTG |
| 32967 | SIVCL757 | 20:07:21 | F | 13.06 | TFP/TFP | FTC/TDF/DTG |
| 33191 | SIVCL757 | 20:07:02 | F | 11.06 | TFP/TFP | FTC/TDF/DTG |
| 34996 | SIVCL757 | 18:06:20 | F | 11.46 | N/A | FTC/TDF/DTG |

### S2 Table. Nonhuman primate cohort for tissue assessment during medculls/necropsies

| Animal ID | Sex | Age (years.months) | Weight (kg) | Infection status | Study Treatments | Euthanasia timepoint at weeks post-infection | Medcull Condition |
|---|---|---|---|---|---|---|---|
| 38163 | F | 15.07 | 7.48 | Uninfected | N/A | N/A | N/A |
| 40691 | M | 12.08 | 12.18 | Uninfected | N/A | N/A | N/A |
| 38691 | F | 14.11 | 13.56 | Uninfected | N/A | N/A | N/A |
| 40499 | F | 12.11 | 9.79 | Uninfected | N/A | N/A | N/A |
| 45721 | M | 5.06 | 14.75 | Uninfected | FTY720 | N/A | N/A |
| 45781 | M | 5.06 | 8.66 | Uninfected | FTY720 | N/A | N/A |
| 46235 | M | 4.07 | 14 | Uninfected | FTY720 | N/A | N/A |
| 46354 | M | 4.06 | 10.96 | Uninfected | FTY720 | N/A | N/A |
| 46410 | M | 4.06 | 12.7 | Uninfected | FTY720 | N/A | N/A |
| 46548 | M | 4.05 | 8.25 | Uninfected | FTY720 | N/A | N/A |
| 46551 | M | 4.05 | 8.59 | Uninfected | FTY720 | N/A | N/A |
| 47081 | F | 3.06 | 5.28 | Uninfected | FTY720 | N/A | N/A |
| 47154 | M | 3.06 | 8.63 | Uninfected | FTY720 | N/A | N/A |
| 47161 | M | 3.06 | 9.32 | Uninfected | FTY720 | N/A | N/A |
| 47387 | F | 3.05 | 7.85 | Uninfected | FTY720 | N/A | N/A |
| 47466 | M | 3.04 | 7.04 | Uninfected | FTY720 | N/A | N/A |
| 34051 | F | 20.00 | 6.97 | Uninfected | N/A | N/A | N/A |
| 35488 | F | 18.01 | 8.54 | Uninfected | N/A | N/A | N/A |
| 36798 | F | 16.11 | 10.31 | Uninfected | N/A | N/A | N/A |
| 37164 | F | 16.01 | 10.83 | Uninfected | N/A | N/A | N/A |
| 40234 | F | 12.11 | 8.72 | Uninfected | N/A | N/A | N/A |
| 40742 | M | 12.00 | 11.81 | Uninfected | N/A | N/A | N/A |
| 41217 | F | 11.11 | 10.77 | Uninfected | N/A | N/A | N/A |
| 41812 | F | 11.00 | 6.51 | Uninfected | N/A | N/A | N/A |
| 25888 | F | 29.01 | 9.92 | Uninfected | N/A | N/A | N/A |
| 33374 | F | 19.08 | 6.64 | Uninfected | N/A | N/A | N/A |
| 35328 | M | 17.03 | 11.22 | Uninfected | N/A | N/A | N/A |
| 47862 | M | 0.08 | 1.27 | Uninfected | N/A | N/A | CHRONIC DIARRHEA |
| 42318 | F | 10.11 | 8.25 | Uninfected | N/A | N/A | NX (UNRELATED PROJECT) |
| 39157 | F | 11.08 | 8.88 | Uninfected | N/A | N/A | BILATERAL RENAL FAILURE |
| 35115 | M | 16.08 | 12.12 | Uninfected | N/A | N/A | BILATERAL ARTHRITIS STIFLES |
| 44003 | F | 5.11 | 5.5 | Uninfected | N/A | N/A | BILATERAL ARTHRITIS STIFLES |
| 44510 | M | 5.09 | 7.47 | Uninfected | N/A | N/A | REGENERATIVE JOINT DISEASE - STIFLES |
| 44288 | F | 5.01 | 4.9 | Uninfected | N/A | N/A | TRICHOBEZOAR; CHRONIC ULCERATIVE GASTRITIS |
| 41647 | M | 8.11 | 9.18 | Uninfected | N/A | N/A | HEPATIC AMYLOID |
| 46078 | F | 19.07 | 6.19 | Uninfected | N/A | N/A | HEMOABDOMEN |
| 33100 | F | 20.07 | 12.32 | Uninfected | N/A | N/A | N/A |
| 33171 | F | 20.07 | 9.27 | Uninfected | N/A | N/A | N/A |
| 33980 | F | 19.07 | 9.6 | Uninfected | N/A | N/A | N/A |
| 33994 | F | 19.06 | 10.31 | Uninfected | N/A | N/A | N/A |
| 34194 | F | 19.06 | 11.84 | Uninfected | N/A | N/A | N/A |
| 35886 | F | 17.06 | 9.11 | SIVCL757 | Embicitrabine, Tenofovir Disproxil Fumarate, Dolutegravir (ART) | 121 | N/A |
| 35595 | F | 17.08 | 11.9 | SIVCL757 | Embicitrabine, Tenofovir Disproxil Fumarate, Dolutegravir (ART) | 121 | N/A |
| 34974 | F | 17.09 | 8.94 | SIVCL757 | Embicitrabine, Tenofovir Disproxil Fumarate, Dolutegravir (ART) | 78 | BICAVITARY EFFUSION |
| 32967 | F | 20.08 | 13.06 | SIVCL757 | Embicitrabine, Tenofovir Disproxil Fumarate, Dolutegravir (ART) | 121 | N/A |
| 33191 | F | 20.07 | 11.06 | SIVCL757 | Embicitrabine, Tenofovir Disproxil Fumarate, Dolutegravir (ART) | 121 | N/A |
| 34996 | F | 18.07 | 11.46 | SIVCL757 | Embicitrabine, Tenofovir Disproxil Fumarate, Dolutegravir (ART) | 121 | N/A |

### S3 Table. Key reagents and resources.

| REAGENT or RESOURCE | SOURCE | IDENTIFIER |
|---|---|---|
| Antibodies | | |
| Mouse anti-Human CXCR3 (Clone:1C6/CXCR3) - APC | BD Biosciences | Cat# 550967; RRID: AB_398481 |
| Mouse anti-Human CD16 (Clone:3G8) - APC | BD Biosciences | Cat# 561248; RRID: AB_10612010 |
| Mouse anti-Human CD49d (Clone: HP2/1) - APC | Beckman Coulter | Cat# B01682; RRID: AB_398681 |
| Mouse anti-Human CD28 (Clone: 28.2) - APC-Cy7 | BioLegend | Cat# 302966; RRID: AB_2800753 |
| Mouse anti-Human CD3 (Clone: SP34-2) - APC-Cy7 | BD Biosciences | Cat# 557757; RRID: AB_396863 |
| Mouse anti-Human CD20 (Clone: 2H7 ) - APC-Cy7 | BioLegend | Cat# 302314; RRID: AB_314262 |
| Mouse anti-Human KI-67 (Clone: B56) - Alexa Fluor 488 | BD Biosciences | Cat#558616; RRID: AB_10611866 |
| Mouse anti- Non-Human Primate CD45 (Clone: D058-1283) - Alexa Fluor 488 | BD Biosciences | Cat# 557803; RRID: AB_396879 |
| Mouse anti-Human BCL2 (Clone: Bcl-2/100) - Alexa Fluor 647 | BD Biosciences | Cat# 563600; RRID: AB_2738306 |
| Mouse anti-Human IL-21 (Clone:3A3-N2.1) - Alexa Fluor 647 | BD Biosciences | Cat# 560493; RRID: AB_1645421 |
| Mouse anti-Human CD3 (Clone: SP34-2) - Alexa Fluor 700 | BD Biosciences | Cat#557917; RRID: AB_396938 |
| Mouse anti-Human CD14 (Clone: M5E2) - Alexa Fluor 700 | BioLegend | Cat# 301822; RRID: AB_493747 |
| Rat anti-Human CCR7 (Clone:3D12) - PE | BD Biosciences | Cat# 552176; RRID: AB_394354 |
| Mouse anti-Human CD163 (Clone: GHI/61) - PE | BioLegend | Cat# 333606; RRID: AB_1134002 |
| Mouse anti-Human IL-21 (Clone:3A3-N2.1) - PE | BD Biosciences | Cat# 562042; RRID: AB_10896123 |
| Mouse anti-Human CD21 (Clone: Bly-4) - PE-Cy7 | BD Biosciences | Cat# 561374; RRID: AB_10681717 |
| Mouse anti-Human CD103 (Clone: Ber-Act8) - PE-Cy7 | BioLegend | Cat# 350212; RRID: AB_10782579 |
| Mouse anti-Human CD11C (Clone: 3.9) - PE-Cy7 | Invitrogen | Cat# 25-0116-42; RRID: AB_1582274 |
| Mouse anti-Human EOMES (Clone: WD1928) - PE-Cy7 | ThermoFisher | Cat# 25-4877-42; RRID: AB_2573456 |
| Mouse anti-Human CD127 (Clone: eBioRDR5) - PE-Cy7 | ThermoFisher | Cat# 25-1278-42; RRID: AB_1659672 |
| Mouse anti-Human IFN-g (Clone: B27) - PE-Cy7 | BioLegend | Cat# 506518; RRID: AB_2123321 |
| Mouse anti-Human CCR6 (Clone: G034E3) - PE-CF594 | BioLegend | Cat# 353430; RRID: AB_2564233 |
| Mouse anti-Human CD40 (Clone: 5C3) - PE-CF594 | BioLegend | Cat# 334342; RRID: AB_2566457 |
| Mouse anti-Human IL-2 (Clone:MQ1-17H12) - PE-CF594 | BioLegend | Cat# 500344; RRID: AB_2564091 |
| Mouse anti-Human TNFA (Clone: MAb11) - FITC | BioLegend | Cat# 502906; RRID: AB_315258 |
| Mouse anti-Human CXCR3 (Clone: 1C6/CXCR3) - BV421 | BD Biosciences | Cat# 562558; RRID: AB_2737653 |
| Mouse anti-Human FOXP3 (Clone: 206D) - BV421 | Biolegend | Cat# 320124; RRID: AB_2565972 |
| LIVE/DEAD Fixable Aqua Dead Cell Stain Kit - BV510 | Life Technologies | Cat# L34966; RRID: N/A |
| Mouse anti-Human CD11b (Clone: ICRF44) - BV510 | BD Biosciences | Cat# 563088; RRID: AB_2737996 |
| Mouse anti-Human CD8 (Clone: SK1) - BV510 | BD Biosciences | Cat# 563919; RRID: AB_2722546 |
| Mouse anti-Non-Human Primate CD45 (Clone: D058-1283) - BV605 | BD Biosciences | Cat# 564098; RRID: AB_2738590 |
| Mouse anti-Human CCR4 (Clone: 1G1) - BV605 | BD Biosciences | Cat# 562906; RRID: AB_2737882 |
| Mouse anti-Human CD4 (Clone: L200) - BV650 | BD Biosciences | Cat# 563737; RRID: AB_2687486 |
| Mouse anti-Human CCR5 (Clone: 3A9) - BV650 | BD Biosciences | Cat# 564999; RRID: AB_2739037 |
| Mouse anti-Human CD69 (Clone: FN50) - BV711 | BioLegend | Cat# 310944; RRID: AB_2566466 |
| Rat anti-Human CX3CR1 (Clone:2A9-1) - BV711 | BioLegend | Cat# 341630; RRID: AB_2814256 |
| Mouse anti-Human CD95 (Clone: DX2) - BUV737 | BD Biosciences | Cat# 564710; RRID: AB_2738907 |

| | | |
|---|---|---|
| Rat anti-Human CX3CR1 (Clone:2A9-1) - BV711 | BioLegend | Cat# 341630; RRID: AB_2814256 |
| Mouse anti-Human CD95 (Clone: DX2) - BUV737 | BD Biosciences | Cat# 564710; RRID: AB_2738907 |
| Armenian Hamster anti-Human/Mouse/Rat ICOS (Clone: C398.4A) - BV785 | BioLegend | Cat# 313534; RRID: AB_2629729 |
| Mouse anti-Human CCR5 (Clone: 3AG) - BV786 | BD Biosciences | Cat# 565001; RRID: AB_2739039 |
| Mouse anti-Human HLA-DR (Clone: L243) - BV786 | BioLegend | Cat# 307642; RRID: AB_2563461 |
| Mouse anti-Human CD8 (Clone: SK1) - BUV805 | BD Biosciences | Cat# 564913; RRID: AB_2833078 |
| Mouse anti-Human PD-1 (Clone: EH12.2H7) - Pacific Blue | BioLegend | Cat# 329916; RRID: AB_2283437 |
| LIVE/DEAD Fixable Near-IR Dead Cell Stain Kit | Life Technologies | Cat# L34976; RRID: N/A |
| Mouse anti-Human CD45 (Clone: 2B11+PD7/26) | Novus | Cat# NBP2-34528AF647; RRID: AB_960384 |
| Rat anti-Human CD3 (Clone: CD3-12) | Bio-Rad | Cat# MCA1477; RRID: AB_321245 |
| Rabbit anti-NeuN (Clone: polyclonal) | Millipore Sigma | Cat# ABN78; RRID: AB_10807945 |
| Goat anti-Rat IgG (H+L) Crossed Adsorbed secondary Antibody, Alexa Fluor 594 | ThermoFisher | Cat# A-11007; RRID: AB_10561522 |
| Syto83 | ThermoFisher | Cat# S11364; RRID: N/A |
| Rabbit anti-Human CD3 (Clone: polyclonal) | Agilent | Cat# A045229-2; RRID: AB_2335677 |
| Mouse anti-Human CD4 (Clone: OTI5D9) | Novus | Cat# NBP2-46149; RRID: N/A |
| Rabbit anti-Human CD11b (polyclonal) | Invitrogen | Cat# PA5-29633; RRID: AB_2547108 |
| Rabbit anti-Human CD45 (Clone: polyclonal) | Abcam | Cat# ab10558; RRID: AB_442810 |
| Rabbit anti-Human IBA1 (Clone: HL22) | Invitrogen | Cat# MA5-36257; RRID: AB_2890455 |
| Rabbit anti-Human NeuN (Clone: EPR12763) | Abcam | Cat# ab177487; RRID: AB_2532109 |
| Purified NA/LE Mouse anti-Human CD49d (Clone: 9F10) | BD Biosciences | Cat# 555501; RRID: AB_2130052 |
| Purified NA/LE Mouse anti-Human CD28 (Clone: CD28.2) | BD Biosciences | Cat# 555725; RRID: AB_396068 |
| **Bacterial and virus strains** | | |
| SIVsm804e-CL757 | Dr. Vanessa Hirsch, NIAID | doi: 10.1371/journal.ppat.1006538 |
| **Biological samples** | | |
| Rhesus macaque biological fluids (Blood and Cerebrospinal Fluid) | CNPRC, UC Davis | N/A |
| Rhesus macaque tissues (Brain tissue, pituitary gland, dura mater, choroid plexus, skull bone marrow, draining cervical lymph node, thoracic lymph node, fine needle axillary lymph node aspirates, spleen) | CNPRC, UC Davis | N/A |
| Formalin fixed and paraffin embedded human hippocampal tissue from non-demented anonymous human patients | Netherlands Brain Bank | N/A |
| Formalin fixed and paraffin embedded human hippocampal tissue from an anonymous human patient with glioblastoma | Netherlands Brain Bank | |
| Formalin Fixed and paraffin embedded human tonsil tissue | Cancer Center Repository, UC Davis | N/A |

| | | |
|---|---|---|
| **Chemicals, peptides, and recombinant proteins** | | |
| Brilliant Stain Buffer plus | BD Biosciences | Cat# 566385 |
| BD FACS Lysing Solution | BD Biosciences | Cat# 349202 |
| BD Cytofix/CytoPerm | BD Biosciences | Cat# 51-2090KZ |
| BD Perm/Wash | BD Biosciences | Cat# 51-2091KZ |
| eBioscience Foxp3/Transcription staining buffer set | Invitrogen | Cat# 00-5523-00 |
| Protein Transport Inhibitor Containing Brefeldin A (BD GolgiPlug) | BD Biosciences | Cat# 555029 |
| Protein Transport Inhibitor Containing Monensin (BD GolgiStop) | BD Biosciences | Cat# 554724 |
| Gill's Hemotoxylin I | StatLab | Cat# HXGH1LT |
| Collagenase Type 4 (265 u/mg dw) | Worthington Biochemical Corporation | Cat# LS004188 |
| DNase I Recombinant, RNase-free | Roche Diagnostics | Cat# 04716728001 |
| Embicitrabine | Gilead Sciences | Cat# GS-9019 |
| Tenofovir Disproxil Fumarate | Gilead Sciences | Cat# GS-4331 |
| Dolutegravir (GSK Comet) | GSK plc | Cat# GSK1349572A |
| Hydroxypropylbetadex (Kleptose HPB) | Roquette | Cat# 346113102B |
| FTY720 | Millipore Sigma | Cat# SMLO700 |
| **Critical commercial assays** | | |
| T cell activation/Expansion Kit, non-human primate | Miltenyi Biotec | Cat# 130-092-919 |
| eBioscience Cell Stimulation Cocktail | Invitrogen | Cat# 00-4970-93 |
| LEGENDplex NHP Inflammation Panel (13-plex) | BioLegend | Cat# 740389 |
| CD45 microbeads, non-human primate | Miltenyi Biotec | Cat# 130-091-899 |
| Chromium Next GEM Single Cell 3' GEM, Library & Gel Bead Kit v3.1 | 10X Genomics | Cat# PN-1000121 |
| Chromium Nuclei Isolation Kit with RNase Inhibitor | 10X Genomics | Cat# PN-1000494 |
| Chromium Next GEM Single Cell Multiome ATAC + Gene Expression Reagent Bundle | 10X Genomics | Cat# PN-1000283 |
| **Deposited data** | | |
| Single Cell RNA Sequencing Results | This paper | GEO Accession: GSE221815 |
| **Experimental models: Organisms/strains** | | |
| Rhesus macaque (Macaca Mulatta) | CNPRC, UC Davis | N/A |
| **Oligonucleotides** | | |
| **Software and algorithms** | | |
| GraphPad Prism (Version 9.5.1) | GraphPad | https://www.graphpad.com/ |
| FACS Diva (Version 8.0.1) | BD Biosciences | https://www.bdbiosciences.com/en-us/products/software/instrument-software |
| FlowJo (Version 10.8.1) | FlowJo LLC | https://www.flowjo.com/ |
| FlowSOM (Version 2.4.0) | Van Gassen et al. (doi: 10.1002/cyto.a.22625.) | https://bioconductor.org/packages/release/bioc/html/FlowSOM.html |
| R (Version 4.2.1) | R Core Team | https://www.r-project.org/ |
| CellRanger (Version 7.0.1) | 10X Genomics | https://support.10xgenomics.com/single-cell-gene-expression/software/pipeli |
| Seurat (Version 4.3.0) | Hao et al. (doi: 10.1016/j.cell.2021.04.048.); Stuart et al. (doi: 10.1016/j.cell.2019.05.031) | https://satijalab.org/seurat/ |
